# Identification and characterization of specific motifs in effector proteins of plant parasites using MOnSTER

**DOI:** 10.1101/2023.07.03.547457

**Authors:** Giulia Calia, Paola Porracciolo, Yongpan Chen, Djampa Kozlowski, Hannes Schuler, Alessandro Cestaro, Michaël Quentin, Bruno Favery, Etienne G.J. Danchin, Silvia Bottini

**Author notes:** Department of Plant Pathology, China Agricultural University, Beijing, China (Yongpan Chen). INRAE, Université Côte d’Azur, CNRS, Institut Sophia Agrobiotech, Sophia-Antipolis, France. contributed as co-first author (Giulia Calia, Paola Porracciolo). contributed as co-last author (Etienne G.J. Danchin, Silvia Bottini). **To whom correspondence should be addressed:** Silvia Bottini, PhD Institut Sophia Agrobiotech UMR 1355 INRAE/7254 CNRS/UniCA 400 route des Chappes – BP 167 06903 Sophia Antipolis Cedex France.

## Abstract

Plant pathogens cause billions of dollars of crop loss every year and are a major threat to global food security. Identifying and characterizing pathogens effectors is crucial towards their improved control. Because of their poor sequence conservation, effector identification in protein sequences predicted from genomes is challenging and current methods generate too many candidates without indication for prioritizing further experimental studies. In most phyla, effectors contain specific sequence motifs which influence their localization and targets in the plant. Although bacterial, fungal and oomycetes effectors have been studied extensively and conserved characteristic motifs have been identified, research on plant-parasitic nematode effectors (PPN) identified some enriched degenerate motifs in only one species so far. The different lifestyles of PPNs might reflect effectors with different functions according to the nematode’s specific needs, thus presenting a high variety of characteristic motifs.

To circumvent these limitations, we have developed MOnSTER a novel tool that identifies <u>clu</u>sters of <u>m</u>otifs of <u>p</u>rotein <u>s</u>equences (CLUMPs). MOnSTER can be fed with motifs identified by *de novo* tools or from databases such as Pfam and InterProScan. The advantage of MOnSTER is the reduction of motif redundancy by clustering them and associating a score. This score encompasses the physicochemical properties of AAs and the motif occurrences. We built up our method to identify discriminant CLUMPs in candidate parasitism proteins of plant-pathogenic oomycetes. We showed the reliability of MOnSTER by identifying five CLUMPs that correspond to the known motifs: RxLR, -dEER and LxLFLAK-HVLVxxP. Consequently, we applied MOnSTER on PPN candidate parasitism proteins and identified peculiar motifs in their sequences. We identified six CLUMPs in about 60% of the known nematode candidate parasitism proteins. Furthermore, we found that specific co-occurrences of at least two CLUMPs are present in PPN candidate parasitism protein sequences bearing protein domains important for invasion and pathogenicity.

The potentiality of this tool goes beyond the candidate parasitism proteins and can be used to easily cluster motifs and calculate the CLUMP-score on any set of protein sequences.

**Authors summary:** Population growth, environmental degradation and climate change are already bringing harm to human communities and the natural world that needs to be addressed rapidly. Ensuring food security for a population that will exceed 9 billion people by 2050 while preserving the environment and biodiversity is a major challenge. Agricultural pathogens, to cause the infection, secrete effector proteins that promote colonization of the host plant. Identifying and characterizing pathogens’ effectors is crucial towards understanding how they manipulate the plant and better combat them. Because of their poor sequence conservation, effector identification in protein sequences predicted from genomes is challenging and current methods generate too many candidates without indication for prioritizing further experimental studies. To address these challenges, we have developed a novel tool called MOnSTER, that identifies and score <u>clu</u>sters of <u>m</u>otifs of <u>p</u>rotein <u>s</u>equences (CLUMPs). MOnSTER is an easy to use tool that can be included in any pipeline needing motif calling and will be of great use to accelerate both computational and experimental studies relating to protein motif discovery. Altogether our findings provide improvements in the understanding of the mechanisms set up by the pathogens to infect the plant and can elucidate important signatures to block the development of plant-pathogen interactions and allow to engineer of durable disease resistance.

## Introduction

Plant pathogens are a major threat to global food security. To cause the infection, pathogenic organisms secrete effector proteins that promote colonization of the host plant by overcoming the physical barriers of plant cell walls, suppressing or evading immune perception, and deriving nutrients from host tissues [1]. Therefore, identifying and characterizing pathogens effectors is crucial towards understanding how they manipulate the plant and better combat them. Effector proteins are often specific to pathogens and essential for causing plant pathology, constituting targets of choice for the development of cleaner and more specific control methods [2]–[4]. Because of their poor sequence conservation, effector identification among the set of predicted proteins from the genome (proteome) is challenging and current methods generate too many candidates without further indication for prioritizing experimental studies. Classically, effector proteins are indirectly identified among the predicted secretome based on the presence of a signal peptide for secretion and a lack of transmembrane region [5], [6]. However, these criteria alone suffer from two main limitations. On one side, the secretome comprises many proteins that are not effectors, on the other side some known effectors do not possess signal peptides for secretion. In most phyla, effectors contain specific sequence motifs which target host proteins with distinct roles in the infection process and control virulence [7]. The best-studied example is effectors secreted via the type III secretion system (T3SS) class of Gram-negative bacterial pathogens which are characterized by a specific motif/domain conferring a repertoire of molecular determinants with important roles during infection [8], [9]. However, these features are not conserved in other bacteria. Indeed, gram-positive pathogens and certain phloem- and xylem-colonizers, such as *Candidatus liberibacter* and *Xylella spp*., do not encode the T3SS. In these bacteria, effector delivery is dependent on the presence of the N-terminal signal peptide, which is required for protein secretion [10]. In fungi, often effectors are small in size and present cysteine-rich sequences [11]. Another well-characterized example is the effectors of the oomycetes pathogens. Oomycetes are eukaryotic filamentous and heterotrophic microorganisms among which, more than 60% of them parasitize plants [12]. Well-known plant pathogens in oomycetes include late blight of potato, sudden oak death, root rot agents (*Phytophthora* species), and downy mildew *Peronospora* and *Bremia* species [13], [14]. These pathogens code for two notable classes of effector proteins RxLR and Crinkler (CRN), that can be predicted by the occurrence of the related motifs, RxLR, -dEER and LxLFLAK-HVLVxxP in the N-terminal region downstream the signal peptide [15]–[17].

Although for some plant pathogens such as oomycetes, effectors have been studied extensively and characteristics motifs have been identified [18], [19], research on Plant-Parasitic Nematode effectors (PPN) did not identify any consensus motif, conserved across multiple species. The most economically important PPNs are the sedentary Root-Knot Nematodes (RKNs) and cyst nematodes [20]. These sedentary parasites induce the formation of a feeding structure that serves as a constant food source for the nematode. Other PPNs are migratory and a whole spectrum of variations exists between endo and ecto parasites, with semi-endoparasites an intermediate between the two extremes [21]. The different lifestyles of PPNs are expected to be reflected in their secretions, which presumably contain effectors with different functions according to the nematode’s specific needs, thus presenting a high variety of characteristic motifs complicating their identification.

A first step toward the identification of motif characteristics of RKN effectors was performed by Vens et al. [22]. The authors developed a bioinformatic tool, called MERCI, to identify motifs with high occurrences in a positive dataset (known effector sequences) and absent in the negative one (non-effector sequences). MERCI uses a graph-based approach incorporating physicochemical features of the amino acids composing protein sequences. By analyzing the known effector sequences of the RKN species *Meloidogyne incognita*, one of the most important known crop pathogens among all [23], they identified 4 motifs. However, at the time of their publication, very few genomes for RKN species were available, and the study was therefore conducted on one single RKN species. Furthermore, the genome used at that time was later shown to be partially incomplete [24]. These limitations prevent the generalization of the previous findings. Da Rocha et al. identified a *cis*-regulatory promoter motif (Mel-DOG box) characteristic of dorsal gland effectors [25]. Recently, Rocha et al. used this motif combined with other criteria to select new putative effectors and validated 14 new dorsal gland-specific candidate effectors expressed in adult females [26]. Although all these studies have contributed to enlarging the list of known effectors, a global characterization of their properties is still missing. Therefore, there is an urgent need for a novel study of the properties of PPN effector sequences and motif research.

By taking advantage of the multitude of proteomes available nowadays for several PPN, we developed a comprehensive motif mining analysis to identify characteristic motifs of candidate parasitism protein sequences of these species. Sequence motifs are usually of constant short size and are often repeated and conserved. Typically, motifs conform to a particular sequence pattern, where certain positions can be constrained to a specific amino acid, whereas others are not [27]. This confers a high degeneration of the motifs yielding a huge list of non-redundant motif sequences and consequently, some motifs that are not characteristics of effector sequences only [28]. Furthermore, different amino acids (AAs) can have similar physicochemical properties, thus different motif sequences can share similar properties. However, most available motif discovery tools do not consider these properties. To circumvent these limitations, we have developed MOnSTER a novel tool that identifies <u>clu</u>sters of <u>m</u>otifs of <u>p</u>rotein <u>s</u>equences (CLUMP) and associates a score to each CLUMP. This score encompasses the physicochemical properties of AAs and the motif occurrences. Overall, one of the key advantages of MOnSTER is that it reduces the redundancy of motifs found by *de novo* tools. Furthermore, already known motifs available in publicly available databases such as Pfam [29] and/or InterProScan [30] can also be used as input of MOnSTER to identify discriminant CLUMPs.

We built up our method to identify discriminant CLUMPs in 1743 candidate parasitism proteins of plant-pathogenic oomycetes. We showed the reliability of MOnSTER by identifying 5 CLUMPs that correspond to the known motifs: RxLR, -dEER and LxLFLAK-HVLVxxP. After this proof of concept, we applied MOnSTER on PPN effector proteins and identified peculiar motifs in their sequences at an unprecedented level. We selected a set of 4395 protein sequences from 13 PPN species belonging to the genera *Meloidogyne, Globodera, Heterodera, Radopholus and Bursaphelenchus*. We identified 6 CLUMPs present in 60% of the known effectors (positive dataset). Of note these CLUMPs were found in only 5% of the sequences of the negative datasets, thus highlighting the enrichment of the identified motifs in effector sequences. Furthermore, we found a specific co-occurrence of at least two CLUMPs in PPN candidate parasitism protein sequences bearing protein domains important for invasion and pathogenicity.

The potentiality of this tool goes behind the candidate parasitism proteins and can be used to easily cluster motifs and calculate the CLUMPs score on any set of protein sequences. Furthermore, we also provide a new scoring system capable of measuring the physicochemical properties of motifs grouped in CLUMPs and a motif alignment algorithm to better explore chemical-physical properties within the CLUMPs. MOnSTER is freely available at https://github.com/paolaporracciolo/MOnSTER_PROMOCA.git.

## Materials and methods

### Datasets

#### Oomycetes

We used proteins from five oomycetes species to create the input datasets for MOnSTER, namely *Phytophthora infestans*, *Phytophthora sojae*, *Phytophthora ramorum*, *Hyaloperonospora arabidopsidis* and *Bremia lactucae*.

##### Positive dataset

The positive dataset consists of 1743 effector proteins belonging to the aforementioned oomycetes obtained from a concatenation of proteins selected from PHI-base database (v4.14) [31], Uniprot (release 2023_02)[32], and the work of Haas et al., (2009) [33], in which they have manually curated the annotations of the proteins. Since the proteins come from different sources, we used CD-HIT (v4.8.1) [34] with the parameters in **Supplementary information,** to filter out identical protein sequences. A total of 1283 proteins are annotated as RxLR effectors, 377 as Crinkler effectors and the last 83 sequences are proteins with no previously identified motif and known to be involved in the host-pathogen interaction.

##### Negative dataset

Proteins in the negative dataset derive all from Uniprot (release 2023_02) and from the oomycetes species cited before filtered from proteins included in the positive dataset and for evident effector-related annotations. Due to the large amount of non-effector proteins remaining from the filtering we firstly used ‘cd-hit’ to reduce protein sequence redundancy and then, to also reduce the unbalance of the final dataset we refined the selection taking only the representative sequences of the orthogroups found with Orthofinder (v2.5.4) [35]. In total 3009 non effector proteins are included in the negative dataset.

##### Motif Discovery

The last input file consists in a list of motifs identified as enriched in the sequences of the positive dataset compared to the sequences of the negative one. We used MERCI and STREME (v5.5.1) [36], with parameters detailed in **Supplementary information**. We imposed different lengths for motifs prediction to be inclusive but more stringent on the motifs in which we are interested. STREME’s output is a list of motifs. Hence, we used the tool FIMO (v5.5.1) [37], with default parameters to extract 246 degenerated motifs from the 4524 different motifs.

We obtained the following numbers of non-redundant motifs: 19 with MERCI and 246 with STREME. Then, we removed the identical motifs and created a single non-redundant list containing all the motifs in the same format, which resulted in 265 different motifs.

#### Plant Parasitic Nematodes (PPNs)

##### Positive dataset

The positive dataset contains candidate parasitism proteins selected to be likely secreted by PPNs in their plant host and belonging to 13 species (*Meloidogyne incognita, Meloidogyne javanica, Meloidogyne arenaria, Meloidogyne hapla, Meloidogyne chitwoodi, Meloidogyne graminicola, Globodera rostochiensis, Globodera pallida, Heterodera havenae, Heterodera glycines, Heterodera schachtii, Radopholus similis, Bursaphelenchus xylophilus)*. We collected candidate parasitism protein from literature mining. More precisely we considered as candidate parasitism protein those proteins for which *in-situ* hybridization experiments showed that the corresponding transcript is present in nematode secretory glands (dorsal or sub-ventral), implying that these proteins are likely secreted by the nematodes into the host plant. The literature mining led to the extraction of 163 proteins from NCBI GeneBank thanks to the NCBI ‘entrez’ API. We also manually extracted 41 sequences from the publications’ core text and Supplementary information. In addition, we downloaded 41 sequences from WormBase ParaSite (www.parasite.wormbase.org, vWBPS17-WS282 [38], [39]), and eight sequences from nematode.net [40]. In total we obtained 229 candidate parasitism protein. We extended the positive dataset with proteins that are non-redundant homologs of the previous candidate parasitism proteins in PPN proteomes. We first used cd-hit-2D with parameters in **Supplementary information**, to cluster sequences from PPNs proteomes and candidate parasitism proteins [41]. We then pooled all the candidate parasitism proteins from closely related *Meloidogyne* species (e.g., *M. incognita*, *M. javanica* and *M. arenaria*) and scanned each corresponding proteome with this multi-species set of sequences using cd-hit. Since the remaining species are genetically distinct, we then scanned each proteome with the relative set of candidate parasitism proteins, except for *H. havenae* and *M. chitwoodi* for which no proteomes were currently available. We merged the two sets of selected candidate parasitism proteins and we performed CD-HIT intra- and inter-species to reduce dataset redundancy (parameters in **Supplementary information**), retaining only sequences having more than 1% divergence and aligning on more than 80% of their length (the longest sequence from each cluster was kept). The final positive dataset includes 546 candidate parasitism proteins from 13 species.

##### Negative dataset

The negative dataset is composed of 3849 protein sequences that we obtained by selecting genes widely conserved across the nematode tree of life and close outgroup species, including many species that are non-parasites. Specifically, we filtered the results from a previous analysis [42] and only retained genes from orthogroups i) conserved in more than 90% (62/64) of the analyzed species including two tardigrade species (outgroups), and ii) presenting less than 10 genes/species/orthogroups to avoid multigenic families, which would lead to overrepresentation of some proteins. To remove the redundancy, we used the same strategy as for the positive dataset (cdhit2D first and then CD-HIT).

##### Motif Discovery

Using the aforementioned software in the same configuration we obtained the following numbers of non-redundant motifs: 40 with MERCI and 229 with STREME applying FIMO. In total, we obtained 269 different motifs.

All datasets are available at https://github.com/paolaporracciolo/MOnSTER_PROMOCA.git and in **Supplementary tables 1.1-1.2 and 2.1-2.2**.

### MOnSTER pipeline

The MOnSTER (MOtifs of cluSTERs) pipeline is composed of three main steps as described in **Figure 1** and in the following paragraphs.

**Figure 1:**
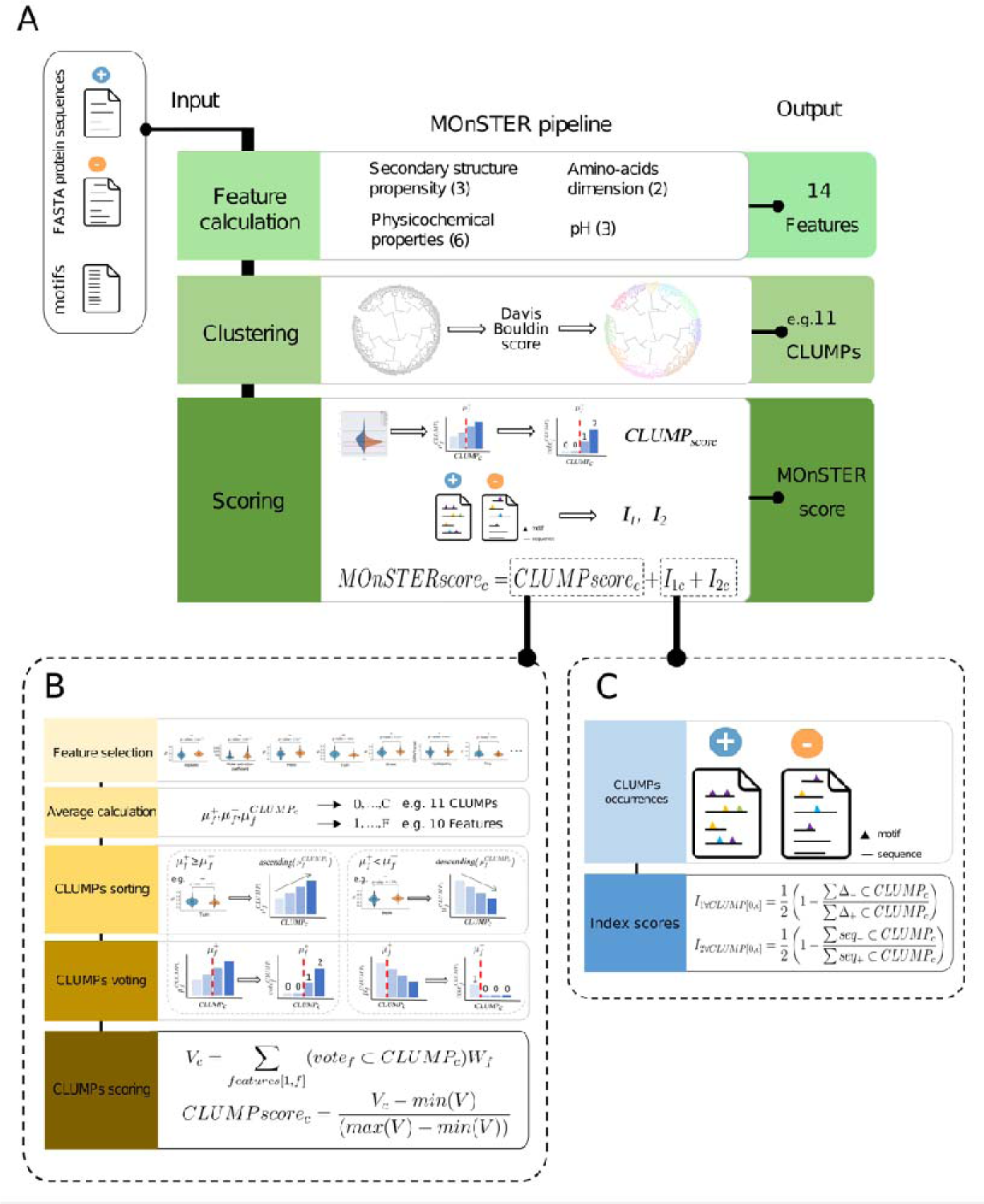
MOnSTER pipeline scheme. (A) MOnSTER pipeline is composed of three steps. It takes two FASTA protein sequences datasets (positive and negative) and a list of predicted motifs (enriched in the positive dataset) as input. The output is a list of CLUMPs and an associated MOnSTER score. The MOnSTER score is constituted by: (B) CLUMP_score_ calculation. (C) Two occurrences Indexes.

#### Feature calculation

The first step of the pipeline concerns the calculation of parameters that describe protein sequences (**Figure 1A**). To allow an easy calculation of the features on any dataset, we calculated the sequence length and used *ProteinAnalysis* class from the *Bio.SeqUtils.ProtParam,* a python sub-package to select 13 additional features based on individual AA properties, belonging to 4 categories:

- secondary structure propensity ‘helix’ (V, I, Y, F, W, L), ‘turn’ (N, P, G, S), and ‘sheet’ (E, M, A, L)).
- amino-acids dimensions (‘tiny’ (A, C, G, S, T) and ‘small’ (A, C, F, G, I, L, M, P, V, W, Y)).
- pH (‘basic’ (H, K, R), ‘acid’ (B, D, E), and ‘charged’ (H, K, R, B, D, E)).
- physicochemical properties (‘hydropathy-score’, ‘polar’ (D, E, H, K, N, Q, R, S, T), ‘non-polar’ (A, C, F, G, I, L, M, P, V, W, Y), ‘aromatic’ (F, H, W, Y), and ‘aliphatic’ (A, I, L, V)).

We performed feature calculations on the positive and negative datasets and the list of motifs. At the end of this step, we obtained three tables of features, one for each of the input datasets (positive, negative datasets and the list of motifs).

#### Clustering

This step allowed to cluster motifs based on their properties described by the 13 features. To make the features comparable to each other, we performed data normalization by using the *StandardScaler* method from *sklearn.preprocessing* [43]. This normalization consists of the removal of the mean and the scaling to unit variance.

Then, we performed a hierarchical clustering of the motifs using the Euclidian distance. We then divided the resulting tree into clusters of motifs of proteins (CLUMPs) selecting the threshold distance that minimized the Davies-Bouldin score [44].

For each CLUMP, we removed the redundant motifs. Briefly, we identified motifs that shared a core sequence (for example: ‘HWT in HWTQ’ and ‘GHWTQ’), and we only retained the cores (for instance: “HWT”) in the CLUMPs.

#### Scoring

The final objective is to identify the CLUMP(s) with the highest discriminative power concerning the positive dataset. Thus, we conceived a new score called the MOnSTER score, to rank the CLUMPs by their discriminative power.

The MOnSTER score is composed of three parts: the CLUMP score and two modified versions of the Jaccard index.

##### CLUMP score

This score considers the AA composition of the motifs belonging to each CLUMP concerning the preferences of the sequences of the positive dataset. The procedure that we implemented to calculate this score is shown in **Figure 1B**.

a) Feature selection

We used the Mann-Whitney test to identify the features whose values were significantly different between the positive and negative datasets. We only retained the statistically significant features, with a p-value < 0.05. Then, we assigned them a score, by calculating -Log(p-value) of each feature. We will refer to it as the ‘feature weight’ hereafter.

b) Average calculation

For each of the selected features (ranging from one to *f*), we calculated the average value for the positive dataset, the negative dataset, and each CLUMP (ranging from zero to *c*). We will refer to these values with the notation: 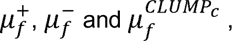 respectively.

c) CLUMPs sorting

We compared the averages of the positive and negative datasets for each feature and sorted CLUMPs accordingly.

Thus, if the 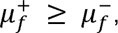 the CLUMPs averages would be sorted in ascending order.

Otherwise 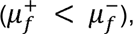 CLUMPs averages would be sorted in descending order.

d) CLUMPs voting

For each feature, and each CLUMP, we divided the CLUMP into two groups accordingly to the following statements:

If 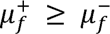: CLUMPs with 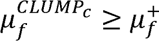 have a vote from 1 to the number of CLUMPs with an increment of 1, otherwise the score is set to 0.

If 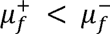: CLUMPs with 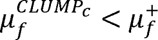 the vote attributed goes from 1 to the number of CLUMPs, otherwise it is 0.

e) CLUMPs scoring

For each CLUMP (ranging from zero to *c*), for each feature (ranging from one to *f*), we multiplied the feature-vote by the ‘feature weight’ (*W_f_*) and summed-up to obtain a CLUMP-vote. Then we scaled each CLUMP-vote to a range from 0 to 1 using the following formula:

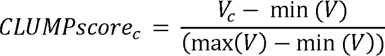

where:

*V* is the list of CLUMPs votes and *V_c_* is calculated as:

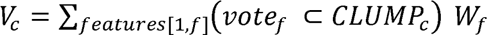

##### Occurrences indexes

The two indexes respectively consider: i) the occurrences of the motifs, for each CLUMP, in the positive dataset compared to the negative, and ii) the number of positive sequences containing the motifs in each CLUMP concerning the negatives (**Figure 1C**).

a) CLUMPs occurrences

We calculated the occurrences of the motifs in each CLUMPs in the two datasets (positive and negative).

b) I’s scores

We propose two ways to calculate the dissimilarity between two sets that will be called I_1_ and I_2_ hereafter.

To obtain I_1_, we calculated the number of occurrences of the motifs for each CLUMP (ranging from zero to *c*) in the negative dataset over the number of occurrences of the motifs of the same CLUMP in the positive dataset, using the following equation:

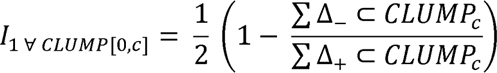

Where:

Δ_-_ and Δ_+_ the number of occurrences of the motifs of the CLUMP in the negative or in the positive dataset, respectively.

To obtain I_2_, for each CLUMP (ranging from zero to *c*), we calculated the number of sequences of the negative dataset that contain at least a motif of the CLUMP, over the number of sequences of the positive dataset that contain at least a motif of the same CLUMP, accordingly to the following formula:

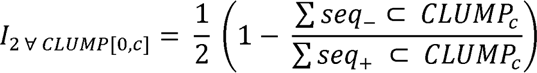

Where:

seq_-_ is the number of sequences of the negative dataset containing at least a motif of the CLUMP.

seq_+_ is the number of sequences of the positive dataset containing at least a motif of the CLUMP.

The ½ factor is applied to have values between 0 and 0.5 for each Index to have equal weight in the final score, and (1 – Index) is to consider the degree of dissimilarity rather than similarity.

##### MOnSTER score

The MOnSTER score, for each CLUMP (from zero to *c*), is the sum of the corresponding CLUMP score, and the two I indexes:

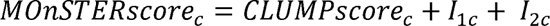

### PRO-MOCA: a novel method to create motif logo of CLUMPs

To create motif logos for each CLUMP, we developed a novel method. PRO-MOCA (PROtein-MOtifs Characteristics Aligner) aligns protein motifs based on the characteristics of the amino acids as shown in **Supplementary figure 1**. The first step is to define the alphabets associated with each characteristic that can be used to represent the motifs (**Supplementary figure 1**a). We have defined four alphabets, namely: “chemical”, “hydrophobicity”, “charge”, “secondary structure propensity” (details for each alphabet are included in the **Supplementary information**).

These alphabets are easily modifiable and other alphabets can be included. Different CLUMP logos can be obtained depending on the alphabet chosen. Secondly, PRO-MOCA uses the selected alphabet to translate the AA sequences of each motif in a CLUMP in the new alphabet (**Supplementary figure 1**b). The third step is the alignment (**Supplementary figure 1**c). Briefly, PRO-MOCA screens the translated motif sequences of a CLUMP looking for a “summit position” with the highest frequency of the same “letter” of the novel alphabet (further details in supplementary materials). Once this position is identified, all motifs are aligned accordingly (**Supplementary figure 1**d). Since the motifs of a CLUMP can have different lengths, after the alignment, PRO-MOCA calculates the number of gaps to add at the extremities to make all motifs having the same length. Importantly, gaps are not allowed inside the motif sequences. The last step of the method is to re-translate the aligned motifs in the original AA sequences (**Supplementary figure 1**e). The alignment is ready to feed a program to create logos. Here we used the tool Weblogo3 [45].

### PPNs candidate parasitism protein domains mining analysis

To investigate the relationship between the selected CLUMPs and functional domains in candidate parasitism proteins we first selected proteins from the positive datasets containing at least one occurrence of a selected CLUMP (311 proteins in total). Then we predicted the protein domains with InterProScan (v5.54-87.0) [30] with default parameters. From the results, we extracted the proteins containing the most frequent predicted domains and considered only entries coming from: MobiDB-lite, Coils, CDD, PANTHER, Pfam and ProSitePatterns. Afterwards, we also predicted the presence of Signal Peptide (SP) (SignalP4.1 [46]) and TransMembrane (TM) domain regions (TMHMM2.0 [47]). We obtained 258 proteins having at least a CLUMP and one of the aforementioned predicted domains, SP or TM.

### *In situ* hybridisation (ISH) and *N. benthamiana* agroinfiltration

*M. incognita* strain “Morelos” was multiplied on tomato (*Solanum lycopersicum* cv. “Saint Pierre”) growing in a growth chamber (25°C, 16 h photoperiod). Freshly hatched J2s were collected and ISH performed as previously described [48], [49]. The *M. incognita Minc3s00056g02931*/*MiEFF72* coding sequence (CDS) lacking the signal peptide for secretion was amplified by PCR with specific primers (EFF72_F: 5’-AAAAAGCAGGCTTCACCATGAATACTGCTGACAAGACACAG-3’ and EFF72_R: 5’-AGAAAGCTGGGTGTTAGAACAAAGCTCGCACTGC-3’) and inserted into the pDON207 entry vector. Antisense probe was amplified using EFF72_R from entry vector. Sense probe was amplified using EFF72_F and used as negative control. Images were obtained with a microscope (Axioplan2, Zeiss, Germany).

The *M. incognita MiEFF72* CDS lacking the signal peptide was recombined in pK7FGW2 (P35S:eGFP-MiEFF72) with Gateway technology (Invitrogen). The construct was sequenced (GATC Biotech) and transferred into *Agrobacterium tumefaciens* strain GV3101. Transient expression was achieved by infiltrating *N. benthamiana* leaves with *A. tumefaciens* GV3101 strain harbouring the GFP-fusion construct, as previously described [50]. Leaves were imaged 48 hours after agroinfiltration, with an inverted confocal microscope (LSM880, Zeiss, Germany) equipped with an Argon ion and HeNe laser as the excitation source. GFP emission was detected selectively with a 505-530 nm band-pass emission filter.

## Results & Discussion

### MOnSTER identified five CLUMPs containing known motifs characteristics of oomycetes effector protein sequences

Characteristic motifs of oomycetes effector proteins are well-known in the literature, such as RxLR, - dEER and LxLFLAK-HVLVxxP [15]. Thus, we reasoned to apply our novel tool, MOnSTER, on oomycetes effectors to test its ability to recover well-characterized motifs. We compiled a set of 4752 oomycetes proteins, comprising 1743 effectors and 3009 non effectors, from five oomycetes species. We performed motif discovery on this set of proteins using MERCI and STREME and we identified 265 significantly enriched motifs (see methods for further details). Then we fed MOnSTER with these motifs and we obtained 11 CLUMPs (**Supplementary table 3**), employing the Davis-Bouldin score, as a criterion to cut the tree. By selecting CLUMPs having a MOnSTER score greater than the median of the overall scores we identified six CLUMPs (CLUMP7, 4, 10, 6, 2 and 9), the first five best-scoring CLUMPs, accordingly to the MOnSTER score, correspond to the known motifs (**Figure 2**). In **Supplementary figure 2** we can also observe that the motifs are respectively grouped in two clades, the two characteristics motifs of CRN-effectors (LxLFLAK and HVLVxxP), form a separate subclade on the right, while the RxLR and -dEER motifs fall into the left clade, resembling the family distinction of effectors to which they belong. More precisely RxLR motifs are divided into two different CLUMPs; CLUMP6 containing only RYLR and RFLR motifs, and CLUMP10, containing other RxLR motifs and included in the same sub-clade of the dEER motif (CLUMP2). The last best-scoring CLUMP contains no known motifs, perhaps suggesting a novel putative motif for oomycetes effectors to investigate. Since oomycetes effectors characterization is not in the scope of this article, we did not consider this last CLUMP for further analysis. In support of that, CLUMPs 7, 4, 10, 6 and 2 are present in 1205/1743 effectors (∼70% of the sequences in the positive dataset) while in combination with the last significant CLUMP (CLUMP9) only two more sequences can be detected.

**Figure 2:**
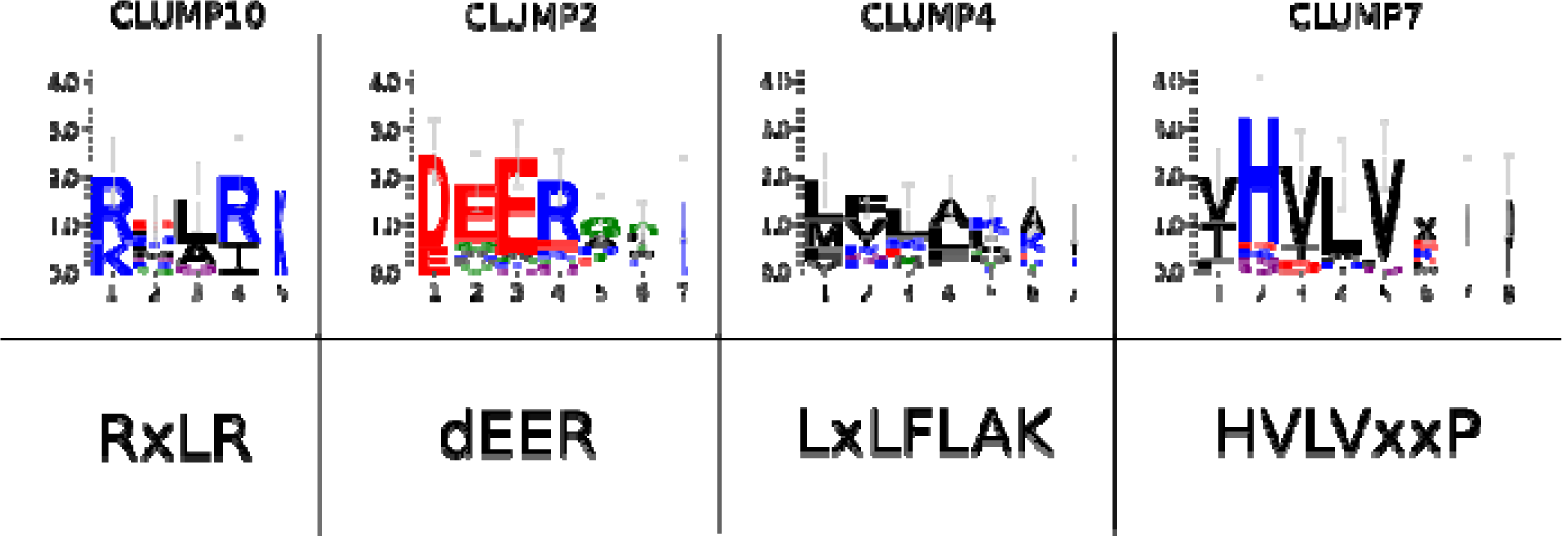
Motif logos of CLUMPs compared to the target motifs. Upper-panel: alignments of motifs in the respective CLUMP are produced by PROMOCA, and then the aligned motif sequences are used to produce the logos with WebLogo3. The x-axis represents the AA position in the motif, while the y-axis represents log-transformed frequencies translated into bits of information. Lower-panel: characteristic motifs of oomycetes effectors families from literature.

Thus, we investigated the occurrences and co-occurrences of the five selected CLUMPs in oomycetes effectors and non-effectors (**Supplementary figure 3**). For the effectors we deeply analyzed the two distinct families; in total we found that 68% of the RxLR-effectors in the positive dataset contain the motifs in CLUMPs associated with the RxLR motif (CLUMP10, 6 and 2). In particular, CLUMP10 and 6 are present alone in 41% of the RxLR-effectors (1238/1743 RxLR-effectors), while 19% of the RxLR-effectors contained the co-occurrence of these CLUMPs with the CLUMPs representing the dEER motif (CLUMP2). This reflects the importance of the RxLR motifs in the effector sequences and the role of the attached dEER [51]. On the other hand, the co-occurrence of CLUMPs specific for LxLFLAK and HVLVxxP (CLUMP7 and 4), in CRN-effector sequences accounts for 67% of the relative sequences in the positive dataset (377/1743). The high co-occurrences rate of CLUMP7 and 4 is strongly in agreement with the presence of LxLFLAK and HVLVxxP motif marking the beginning and the end of the DWL-domain in the Crinkler-effector family [33]. For the negative dataset, instead, only 15% of the sequences show the presence of CLUMP-motifs with a huge decrease in CLUMPs co-occurrences. Overall co-occurrences, indeed, are present in around 30% of positive sequences and in 1% of negative ones.

Previous research showed that the motifs characteristics of oomycetes effectors have strong sequence position preferences [52]–[54]. Thus, we plotted the CLUMPs occurrences in the positive versus negative dataset (**Supplementary figure 4**). Indeed, we can observe that the CLUMPs are concentrated at the beginning of the sequence in positive sequences and conversely spread around the sequence of negative dataset proteins. More precisely the five most interesting CLUMPs are condensed in the first 40% of the sequence with a higher preference at the very beginning and around 30% of the sequence probably corresponding to the N-terminal of the protein in which the target motifs lie.

Altogether these results highlight the ability of MOnSTER to identify CLUMPs containing biologically relevant motifs.

### MOnSTER allowed to identify six CLUMPs characteristics of nematode candidate parasitism proteins

The application of MOnSTER of the oomycetes effectors served as a proof of concept of our methodology. Thus, we moved to the characterization of nematode candidate parasitism sequences for which no characteristic motifs have been identified yet. We collected a set of 4395 proteins, including 546 well-known candidate parasitism proteins and 3849 proteins in the negative dataset, coming from 13 nematode species. By running motif discovery analysis as for the previous dataset, we found 269 motifs enriched in the candidate parasitism protein sequences. By applying MOnSTER with the previous configuration, the 269 input motifs were grouped into 11 CLUMPs. Six best-scoring CLUMPs were selected using the median as the significant threshold (**Supplementary table 4**). Similar to the oomycetes results, we observe two main clades (**Figure 3**): the second and the third best scoring ones (CLUMP2 and 5 respectively) form a single clade while the other significant CLUMPs (CLUMP1, 3, 7 and 10) are distributed in the bigger clade with the non-significant ones. Overall, we found at least one occurrence of one of the six CLUMPs in almost 60% of sequences from the positive dataset compared to 5% of sequences from the negative.

**Figure 3:**
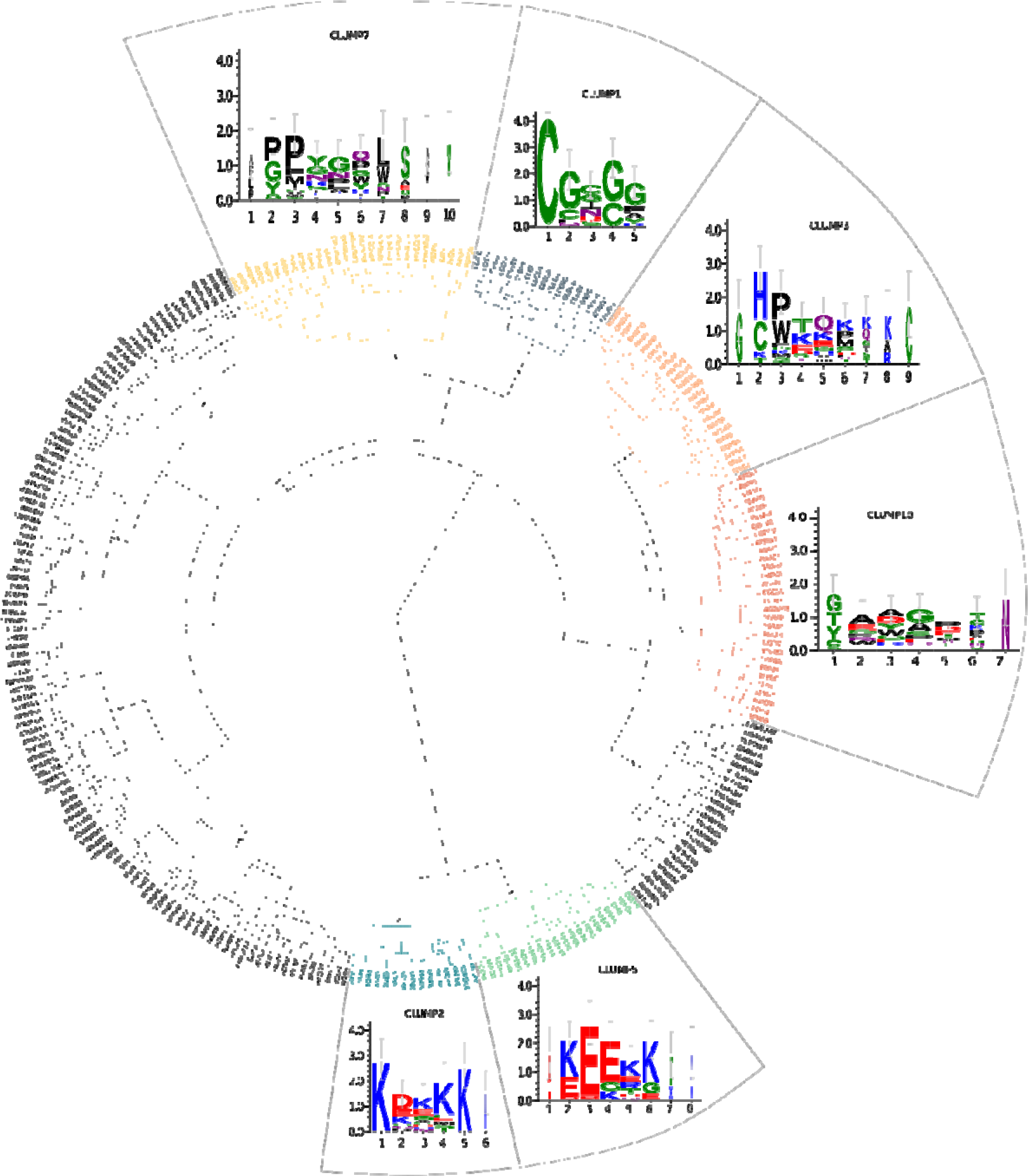
Dendrogram of CLUMPs in Plant Parasitic Nematodes (PPNs) 11 CLUMPs produced by MOnSTER (indicated with “/” sign). The coloured ones are those selected as best-scoring CLUMPs after MOnSTER-score calculation. Each best-scoring CLUMP is associated with the corresponding motif logo; alignment of motifs in each CLUMP is produced by PROMOCA and then WebLogo 3 is used to produce the image (the x-axis shows the AA position of the motif and the y-axis represents the log-transformed frequency of each AA in terms of bits of information).

Then we investigated the presence of the six CLUMPs in each of the 13 PPN species present in the dataset. **Figure 4** shows the abundance of the six best-scoring CLUMPs in the species according to their phylogeny tree. The first three species are the most represented in the positive dataset. Interestingly very distant species show similar CLUMPs frequencies thus suggesting that they might share common characteristics at the sequence level for accomplishing similar functions. Furthermore, we could identify characteristic CLUMPs also for species represented in the dataset with very few sequences reinforcing the previous observation. Overall, this analysis suggests that CLUMPs might be associated with the functional properties of PPN nematodes.

**Figure 4:**
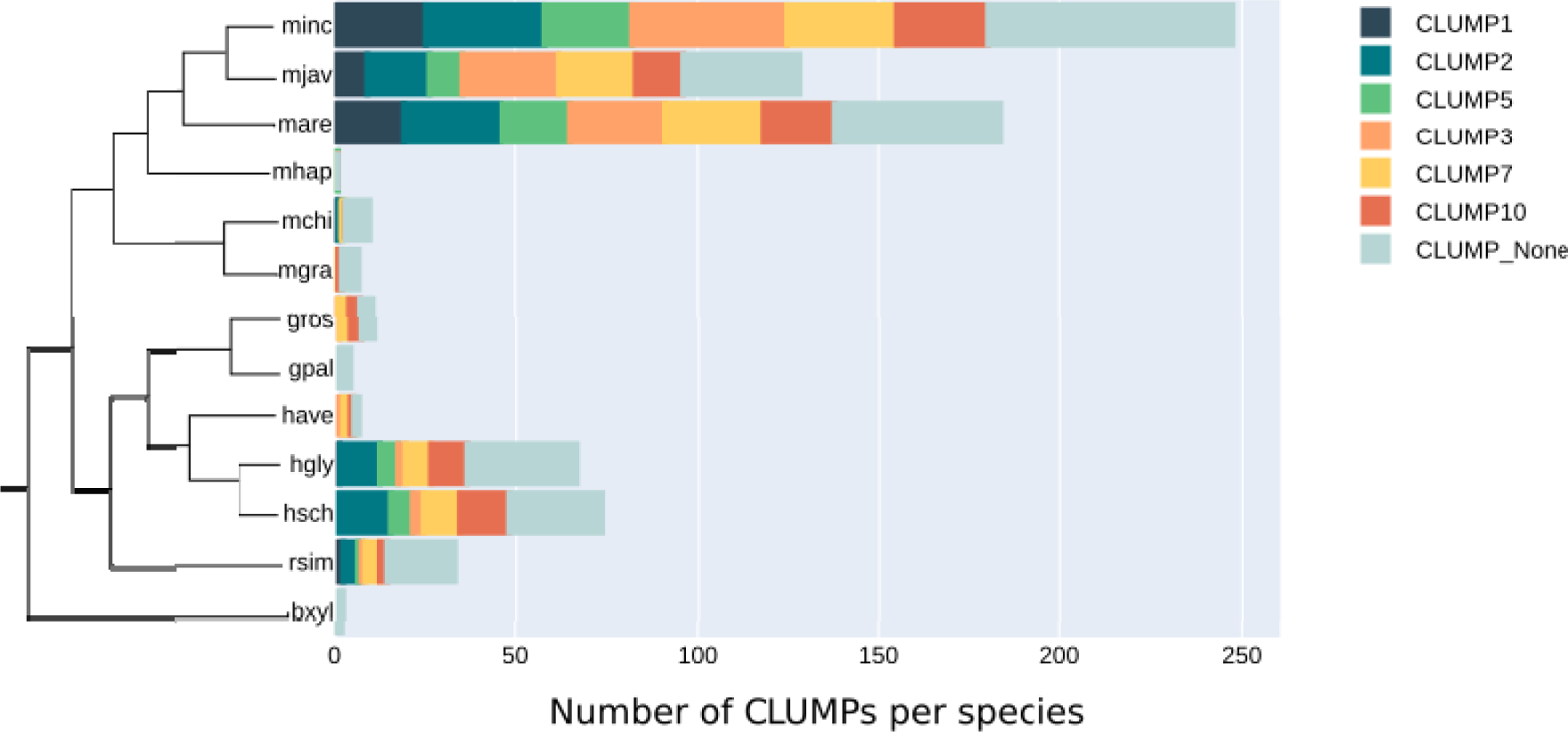
Cardinality of CLUMPs-motifs in each PPN species considered. The total number of motifs belonging to each significant CLUMP per PPN species accordingly to their phylogeny. (minc: *Meloidogyne incognita,* mjav: *Meloidogyne javanica,* mare: *Meloidogyne arenaria,* mhap: *Meloidogyne hapla,* mchi: *Meloidogyne chitwoodi,* mgra: *Meloidogyne graminicola,* gros: *Globodera rostochiensis,* gpal: *Globodera pallida,* have: *Heterodera havenae,* hgly: *Heterodera glycines,* hsch: *Heterodera schachtii,* rsim: *Radopholus similis,* bxyl: *Bursaphelenchus xylophilus*)

Finally, we focused on the positional sequence preferences of CLUMPs in candidate parasitism protein sequences (**Supplementary figure 5**). In general, we observe a difference in the position preferences of the best-scoring CLUMPs between positive and negative dataset sequences. The six CLUMPs tend to occur more frequently in the middle of the sequences in candidate parasitism proteins (positive dataset), with more abundance in central (around 50% of the sequence) and terminal (around 70%), positions. The same CLUMPs are rare in the central position of the negative dataset protein sequences (negative dataset). Contrary to the properties of oomycetes effectors, whose characteristics CLUMPs occur mainly at the beginning of the sequence, PPN candidate parasitism proteins showed a different pattern of occurrences, privileging a central – C terminal occurrence.

#### Co-occurrences of different CLUMPs are associated with functional protein domains

We investigated the co-occurrence patterns of CLUMPs in the PPNs candidate parasitism protein sequences (all possible combinations of co-occurrences are reported in **Supplementary figure 6)**. Overall, we notice that CLUMPs tend to co-occur more frequently in the sequences of the positive dataset than in the negative one, despite the positive set being smaller than the negative one. 30% of candidate parasitism protein sequences show co-occurrences of the six selected CLUMPs, while in the sequences from the negative dataset, co-occurrences, are present in less than 1% of the sequences. As observed for oomycetes, some CLUMPs tend to be present alone, while others tend to co-occur with specific CLUMPs. This suggests that different classes of nematode candidate parasitism proteins might exist, similar to the oomycetes effectors. Interestingly, among the 311 candidate parasitism proteins bearing at least one occurrence of one of the six selected CLUMPs, 72 do not have a predicted signal peptide, consisting of 55% of the proteins in the positive dataset not having the signal peptide. Of note, this is a similar percentage to the percentage of proteins bearing both the CLUMPs and the signal peptide, suggesting that CLUMPs characterize sequence properties beyond the type of secretion. Furthermore, similar patterns of co-occurrences of CLUMPs in candidate parasitism proteins bearing or not the signal peptide are observed with slightly higher co-occurrence presences in the sequences not having the signal peptide (**Supplementary figure 7**). Importantly, there is no relationship between the sequence length and the number of co-occurrences possibly suggesting a functional role for CLUMPs co-occurrences (**Supplementary figure 8**).

To inspect further a putative functional role of CLUMPs in candidate parasitism protein sequences, we queried the sequences having at least one CLUMP or a co-occurrence of multiple CLUMPs against several protein domain databases (see Methods, results in **Figure 5** and **Supplementary table 5**). Among the 311 candidate parasitism protein sequences bearing at least one occurrence of at least one of the six CLUMPs, 84 also have at least an occurrence of a known protein domain. The most recurrent hits are the coil domain, intrinsically disordered domain and the presence of the signal peptide (SP) followed by the pectate lyase domain, glycosyl hydrolase family 5, Stichodactyla toxin (ShK) domain, 14-3-3 family and cysteine-rich domain. Importantly, none of these domains was also found in the sequences from the negative dataset bearing at least one occurrence of at least one of the six CLUMPs. Interestingly, we observe the almost exclusive association between CLUMPs and functional domains, mainly when multiple CLUMPs co-occur in candidate parasitism protein sequences.

**Figure 5:**
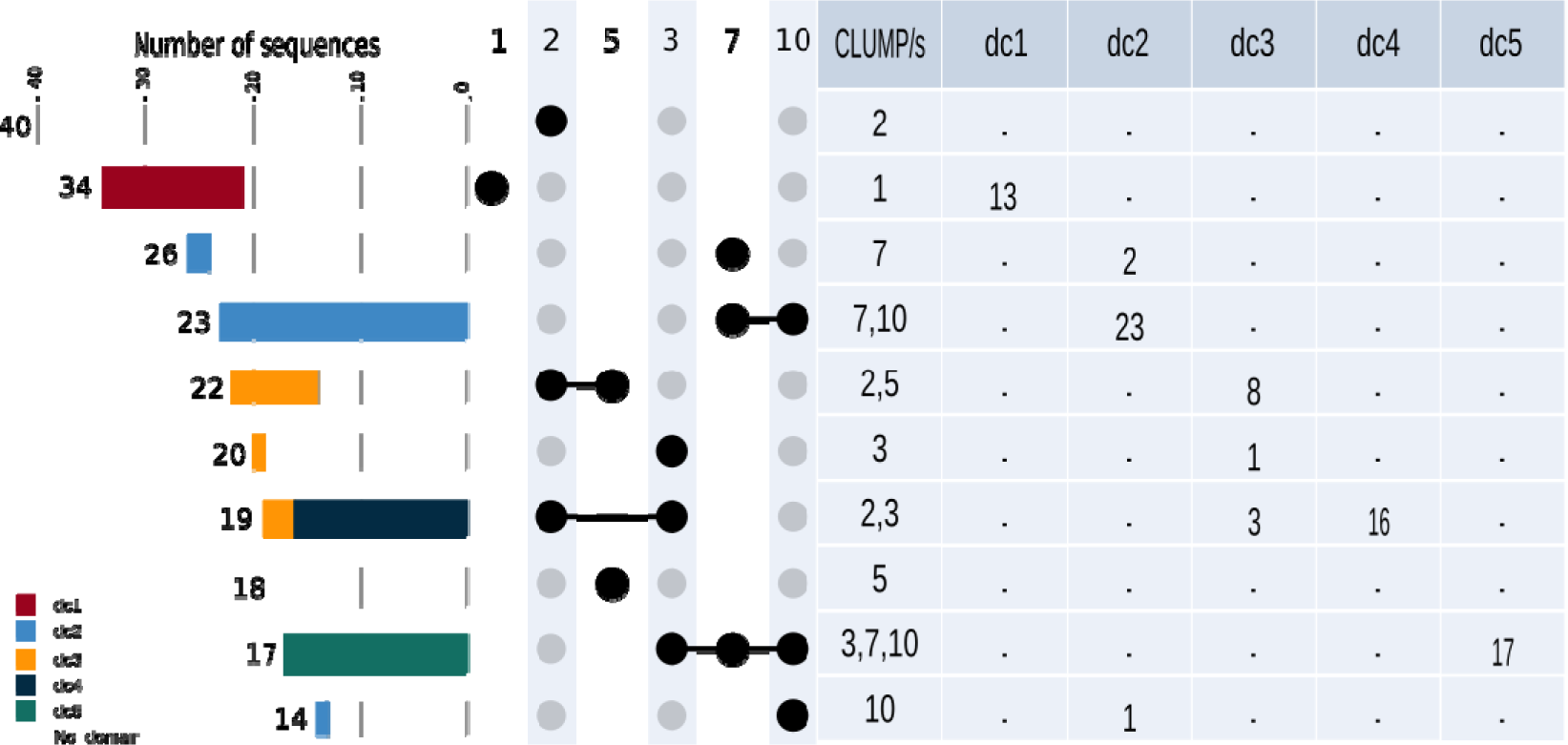
Candidate parasitism proteins showing the presence of CLUMP/s associated with pathogenicity-related protein domain/s. The table on the right shows the co-occurrence of CLUMP or CLUMPs with specific domain classes (dc); dc1, pectate lyase domain class, dc2, glycosyl hydrolase family 5 domain class, dc3 Stichodactyla toxin (ShK) domain class, dc4 14-3-3 family domain class and dc5, cysteine-rich domain class. The upset plot on the left represents the occurrences and co-occurrences of respective CLUMPs in the positive dataset, highlighting the sequences that also have an interesting protein domain following the table counts.

The strongest association that we observe is between the co-occurrences of CLUMPs 7 and 10 and the glycosyl hydrolase family 5 domain on one hand and the co-occurrences of CLUMPs 3, 7, 10 and the cysteine-rich domain, on the other hand. Specifically, all 23 candidate parasitism protein sequences containing the co-occurrences of CLUMP 7 and 10 bear also the glycosyl hydrolase family 5 domain. By inspecting the position of CLUMPs occurrences within the sequences, we observed that the two CLUMPs are flanking the domain: CLUMP7 is consistently present at the beginning of the sequence and consequently of the domain, while CLUMP10 mostly concentrates at the end of the domain, around 60-80% of the sequences (**Supplementary figure 9**). Examples of these genes in nematodes is poorly characterized and likely resulting from horizontal transfer [55], [56]. Similarly, all 17 sequences presenting the co-occurrence of CLUMPs 3, 7,10 also contain the cysteine-rich domain. Cysteine-rich domain and CAP protein are known to be involved in the virulence of nematodes [57]. They are expressed in both plants and pathogens; in the latter, they are important for their virulence by suppressing the host’s immune responses and promoting colonization. Interestingly, these sequences do not contain disordered regions or coil domains, consistently with unique conserved sandwich fold with a large central cavity of these kinds of proteins [58]. 16 out of 19 sequences presenting co-occurrences of CLUMPs 2, 3 have also the 14-3-3 family domain, a eukaryotic-specific protein family with a general role in the signal transduction [59]. We also observe only one motif from CLUMP 2 in these sequences (KDKM) and 4 from CLUMP 3 (NKDKAC, KMKG, PTHPIR, PTHP). 13 out of 34 sequences bearing only CLUMP 1 also contain the pectate lyase domain. Of note, these sequences do not contain coiled or disordinate regions, and only seven show the presence of the SP. Pectate lyase enzymes in nematodes facilitate penetration in plant-cell walls made of pectin [60]. Numerous recent reports showed that these enzymes are produced in specialized nematode gland cells and secreted during the parasitism process. In the case of sedentary endo-parasitic nematodes, this occurs mainly during juvenile migration through the root tissue, when these enzymes play a crucial role in the maceration of the plant tissue facilitating the infection [61]. Finally, eight out of 22 sequences bear the co-occurrences of CLUMPs 2, 5 and the ShK domain. Although the exact biological function of the ShK domain remains unclear, previous reports have shown that this domain might be associated with immunosuppression [62], [63].

Overall, these findings highlight that specific CLUMPs co-occurrences are associated with specific functional domains with roles in invasion and/or infection and might suggest different classes of candidate parasitism proteins cross-species.

#### CLUMPs screening yielded the identification of a novel effector in *M. incognita* validated by *in situ* hybridization

To inspect whether the novel-identified CLUMPs could also help to find new effectors, we focused on the selection of a novel putative effector to validate experimentally. Thus, we selected all proteins of *Meloidogyne incognita* proteome bearing the signal peptide for secreted proteins and no transmembrane domain. Then we screened these sequences and retrieved the ones containing at least one motif of the six significant CLUMPs. Among them, 23% contain at least one occurrence of motifs in CLUMP5 (Supplementary Table 6). Since this is the most abundant CLUMP in this species, we decided to focus on this one to identify a putative candidate to validate experimentally. By literature mining, we refined our list, by sorting out seven sequences that were already experimentally validated by previous studies (Supplementary Table 6). Then we filtered out any candidates having homologs in species other than root-knot nematodes and more than two gene copies to avoid dealing with multigene families according to [42]. Finally, among these eight new putative effector sequences, we studied the pattern of expression of one candidate: *MiEFF72* (*Minc3s00056g02931*) by performing *in situ* hybridisation (ISH). A specific signal was detected in the subventral oesophageal gland cells of pre-parasitic J2s after hybridisation with digoxigenin-labelled MiEFF72 antisense probes (**Figure 6A**). No signal was detected in pre-J2s with sense negative controls. MiEFF72 fused to the C-terminus of GFP was transiently expressed in *N. benthamiana* leaf epidermis. GFP fluorescence was detected in the cytoplasm and in cytoplasmic vesicles (**Figure 6B**). This finding suggests that MiEFF72 be secreted and play a role *in planta* in nematode parasitism.

**Figure 6:**
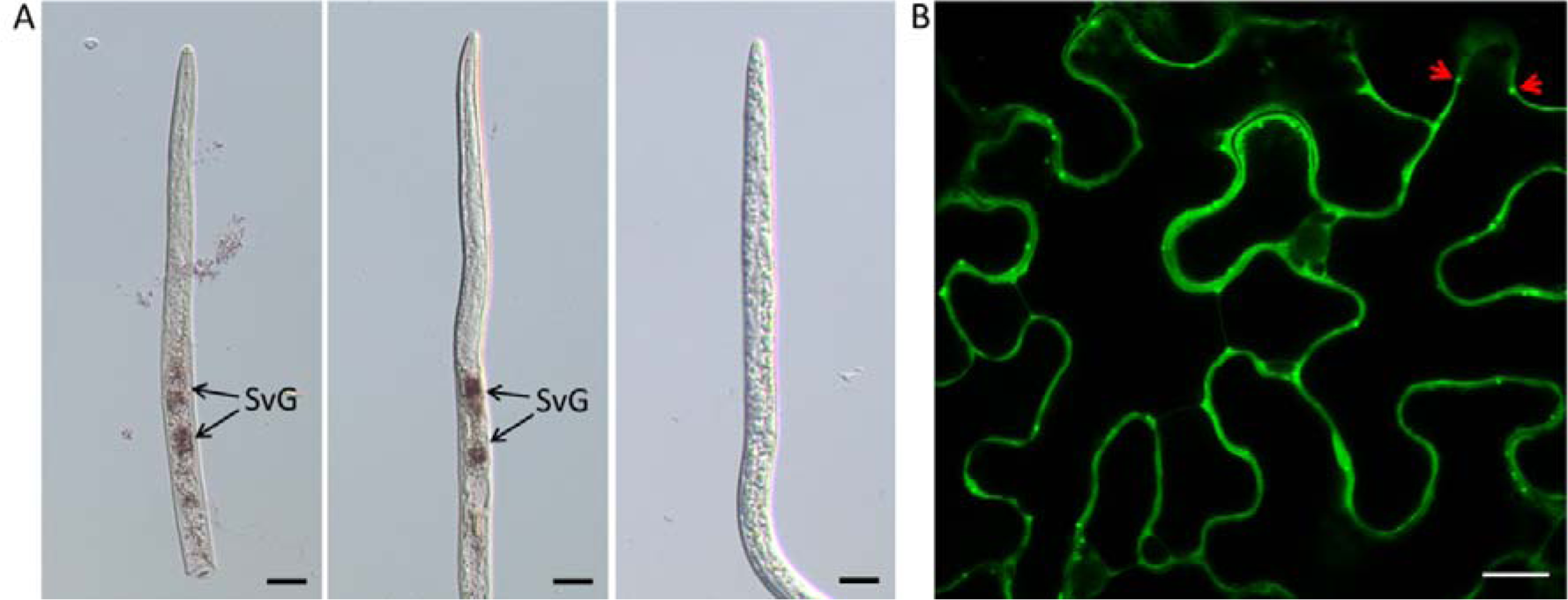
MiEFF72 is specifically expressed in the subventral glands. (A) *In situ* hybridisation showing EFF72 transcripts in the subventral glands (SvG) of J2s of *M. incognita* (two left pictures). Sense probe for the MiEFF72 transcripts was used as a negative control (right picture). (B) MiEFF72 localised to the cytoplasm of plant cells and in cytoplasmic vesicles (red arrows). The MiEFF72 sequence was fused to C-terminal end of the GFP and expressed in *N. benthamiana* leaves by agroinfiltration. Bars = 20 µm.

## Conclusions

This work is structured around three main aims: (1) the development of a novel method to cluster and score discriminant motifs of protein sequences called MOnSTER, (2) the validation of the MOnSTER results by applying it to identify CLUMPs specific to candidate parasitism protein sequences of oomycetes (3) the application of MOnSTER to protein sequences from plant-parasitic nematodes with unprecedented discriminant motifs detection.

The application of MOnSTER on oomycetes yielded the identification of five CLUMPs corresponding to the well-known effector-related motifs like RxLR-dEER and LxLFLAK-HVLVxxP motifs in oomycetes. This demonstrated that the novel scoring method introduced by MOnSTER is a good parameter with which calculate CLUMP specificity for effector protein sequences. When applied to the nematodes candidate parasitism protein, MOnSTER found six novel CLUMPs, not previously characterized. The main advantage of MOnSTER is that the definition of CLUMPs allowed us to reduce the degeneration of 265 and 269 motifs (oomycetes and nematodes respectively), to 11 CLUMPs. Candidate parasitism protein sequences of both pathogens show some common characteristics. Indeed, selected CLUMPs-motifs are present in about 70% of the input proteins for oomycetes and 60% in PPN compared to 15% and 5% in the negative dataset proteins, respectively. Furthermore, around 30% of candidate parasitism protein sequences have co-occurring CLUMPs, in contrast with less than 1% of the negative dataset sequences, in both applications. The main difference between candidate parasitism protein specific motifs of the two pathogens is the positional preference: the beginning of the sequence for oomycetes and central C-terminal for PPNs. This highlights MOnSTER ability to cluster motifs specifically relevant for candidate parasitism protein sequences without privileging any portion of the sequence, like other motif discovery tools.

Concerning the novel identified motifs for PPNs candidate parasitism proteins, we observed that the pattern of occurrences and co-occurrences of CLUMPs in candidate parasitism protein sequences is associated with specific functional domains and might suggest the existence of different classes of candidate parasitism proteins. Importantly we did not observe any species-related preferences thus implying the generality of these results.

In conclusion, MOnSTER quantifies the motifs and sequence properties in each dataset provided, thus allowing a wide application to other protein classes. Since the MOnSTER score considers the physicochemical properties and occurrences of motifs in CLUMPs concerning the protein sequences provided, it works without the need for a reference dataset. Furthermore, the MOnSTER scores are normalized values, therefore, allowing direct comparison between different studies.

Our results highlighted that MOnSTER is a powerful new method to cluster and score discriminant motifs in protein sequences according to their physicochemical properties and pattern of occurrences. It is also a tool that can be easily used on any set of protein sequences and a list of motifs. Therefore, by constructing a dataset of positive candidate parasitism protein sequences and a negative dataset, MOnSTER can be also used to identify CLUMPs characteristics of fungal or bacterial candidate parasitism proteins. As such, MOnSTER can be included in any pipeline needing motif calling and will be of great use to accelerate both computational and experimental studies relating to protein motif discovery.

## Data availability

The source code and related data are available at: https://github.com/paolaporracciolo/MOnSTER_PROMOCA.git

## Acknowledgements

Microscopy work was performed at the SPIBOC imaging facility of Institut Sophia Agrobiotech, and we thank Dr Olivier Pierre for is availability.

## CRediT author contribution

GC: Methodology, Software, Validation, Formal analysis, Writing – original draft, Visualization. PP: Methodology, Software, Writing – original draft. JC: Investigation, Resources. DK: Software, Resources. HS: Writing – review & editing, Supervision. AC: Writing – review & editing, Supervision. MQ: Investigation, Resources, Writing – review & editing, Supervision. BF: Investigation, Resources, Writing – review & editing, Supervision. EGJD: Conceptualization, Methodology, Writing – review & editing, Supervision. SB: Conceptualization, Methodology, Writing – original draft, Writing – review & editing, Supervision, Project administration.

## Supporting information files legend

**Supplementary figure 1: PRO-MOCA algorithm.**

**(a) DEFINITION OF THE ALPHABET.** The user defines the alphabet to use to translate the protein sequences into patterns. Alphabet are constructed as pairs of keys, a letter defined by the user, and values, a list of amino acids to include in the group. Amino acids from the group will be replaced by the key in the sequence. PRO-MOCA already include the following alphabets. Chemical alphabet: ‘P’ = ‘polar’, ‘H’ = hydrophobic, ‘N’= ‘neutral’, ‘A’ = ‘acid’, ‘B’ = ‘basic’. Hydrophobicity alphabet: ‘P’ = ‘hydrophilic’, ‘N’ = ‘neutral’, ‘B’ = ‘hydrophobic’. Charge alphabet: ‘P’ = ‘positive’, ‘N’ = ‘negative’, ‘U’ = ‘neutral’. Secondary structure propensity alphabet: ‘H’ = ‘helix (α)’, ‘S’ = ‘sheet (β)’, ‘T’ = ‘turn’.

**(b) TRANSLATION.** Using the chosen alphabet (step (a)), the motifs in the CLUMP are translated from the protein alphabet (amino acids) into patterns of the given alphabet. For instance, using the ‘chemical’ alphabet, the ‘GHWT’ motif is translated into ‘PBHP’.

**(c) ALIGNMENT.** (c1) PRO-MOCA: (1) takes the list of patterns, (2) selects the first position of each pattern, (3) calculates how many times each letter is repeated at that position and finds the most repeated letter (PRO-MOCA stores this result at each iteration) (4) After that, patterns with the most repeated letter at the first position, lose that letter, and the rest of the pattern is shifted of -1 position. (5) The result is a list of patterns where part of the patterns has a new first position. From this point, the process (3)-(5) is repeated until (6) the result is a list where at least one pattern does not have any more letters. Until there is a pattern with no more letters (golden rectangle). (c2) Once the first part of the alignment is done and PRO-MOCA gets to (6), it finds the central position of the alignment. To do that, PRO-MOCA stored in (3) the number of repetitions of the most repeated letter. In (c2) PRO-MOCA calculates which is the highest value, and gets the corresponding list of patterns with those first positions.

(d) **GAP INSERTION**. The gap insertion is only allowed at the extremities of the patterns and allows to have an alignment where all the patterns have the same length. To do that, PRO-MOCA calculates, for each pattern, how many letters there are before (xi) and after (yi) the central position and calculates the maximum (m and n) of these values. After that, it subtracts m – xi and n – yi. The result of the subtraction is the number of gaps each pattern has before and after the central position.

**(e) RE-TRANSLATION.** In this step, PRO-MOCA re-translates the aligned patterns into aligned motifs

**Supplementary figure 2: Dendrogram of CLUMPs in Oomycetes.**

11 CLUMPs produced by MOnSTER (indicated with “/” sign). The coloured ones are those selected as best-scoring CLUMPs after MOnSTER-score calculation. Each best-scoring CLUMP is associated the motif logo; the alignment of the motif in each CLUMP is produced by PROMOCA and then WebLogo 3 is used to produce the image (the x-axis shows the AA position of the motif and the y-axis represents the log-transformed frequency of each AA in terms of bits of information).

**Supplementary figure 3: Occurrences and co-occurrences of CLUMPs in positive and negative dataset – Oomycetes.**

Each upset-plot represents the number of sequences in the respective dataset according to the presence of one or more CLUMPs ordered by MOnSTER-score. **(a)** positive dataset, **(b)** negative dataset, (c-d) specifically show the occurrence and co-occurrence of CLUMPs corresponding to RxLR and dEER (CLUMP10, 6, 2), and LxLFLAK-HVLVxxP (CLUMP7,4), in RxLR and Crinkler-effector sequences, respectively.

**Supplementary figure 4: Sequence position preference of motifs in CLUMPs – Oomycetes.**

**(a)** Shows the overall preference position of CLUMPs in the positive (blu trace) compared to the negative (orange trace) dataset. In the rug-plot each line corresponds to a CLUMP-motif position occurrence. **(b)** represents the same distribution in the positive dataset but divided by each CLUMP abundance in the sequence bin. **(c)** shows the same concept but in the negative dataset; **(c.1)** is a zoom-in of the original plot to better visualize the occurrences of CLUMPs. The distributions are all represented in bins of 10% each as absolute positions on the sequence.

**Supplementary figure 5: Sequence position preference of motifs in CLUMPs – PPNs.**

**(a)** Shows the overall preference of CLUMPs in the positive (blu trace) compared to the negative (orange trace) dataset. In the rug-plot each line corresponds to a CLUMP-motif position occurrence. **(b)** represents the same distribution in the positive dataset but divided by each CLUMP abundance in the sequence bin. **(c)** shows the same concept but in the negative dataset; **(c.1)** is a zoom-in of the original plot to better visualize the occurrences of CLUMPs. The distributions are all represented in bins of 10% each as absolute positions on the sequence.

**Supplementary figure 6: Occurrence and co-occurrence of CLUMPs in PPNs.**

Representation of occurrence and co-occurrence of each best-scoring CLUMP in protein sequences, according to their MOnSTER-score. (a) shows the number of sequences and the occurrence of CLUMPs in the positive dataset, while (b) represents the same measures in the negative dataset.

**Supplementary figure 7. Occurrence and co-occurrence of CLUMPs in proteins with or without the signal peptide.**

Representation of occurrence and co-occurrence of each best-scoring CLUMP in protein sequences, accordingly to their MOnSTER-score. (a) shows the number of sequences and the occurrence of CLUMPs in the sequences bearing the signal peptide, while (b) represents the same measures in the sequences without the signal peptide.

**Supplementary figure 8: Relation between CLUMPs abundance and protein sequence length – PPNs.**

**(a)** Shows a non-linear relationship between the occurrences of CLUMPs and the length of sequences in the positive dataset, while **(b)** represents the same but in the negative dataset.

**Supplementary figure 9: Protein sequence position of motifs in CLUMP7 and 10 associated with glycosyl hydrolase family 5 domain (dc2).**

**(a)** CLUMP7 and CLUMP10 motifs-position and respective occurrence along selected protein sequences divided into bins representing 10% sequence portion. (**b**) rug-plot representing the start and end position of the domains (PS00659-ProSitePatterns and PF00150-Pfam) belonging to the class 2 of domains, namely glycosyl hydrolase family 5 domains.

**Supplementary table 1.1: Oomycetes’ positive dataset.**

Detailed description of the protein sequences in the oomycetes’ positive dataset including species, retrieving sources, and corresponding amino acid sequence.

**Supplementary table 1.2: Oomycetes’ negative dataset.**

Detailed description of the protein sequences in the oomycetes’ negative dataset including species, retrieving sources, and corresponding amino acid sequence.

**Supplementary table 2.1: PPNs’ positive dataset.**

Detailed description of the protein sequences in the PPNs’ positive dataset including species, retrieving sources, and corresponding amino acid sequence.

**Supplementary table 2.2: PPNs’ negative dataset.**

Detailed description of the protein sequences in the PPNs’ negative dataset including species, retrieving sources, and corresponding amino acid sequence.

**Supplementary table 3: Oomycetes’ MOnSTER-score.**

Summary table of MOnSTER-score for each CLUMP found in oomycetes dataset, ranked by higher score.

**Supplementary table 4: PPNs’ MOnSTER-score.**

Summary table of MOnSTER-score for each CLUMP found in PPNs dataset, ranked by higher score.

**Supplementary table 5: InterProScan domains and CLUMPs co-occurrences.**

Summary of co-occurrences of InterProScan domains in the six characteristic CLUMPs for PPN species.

**Supplementary table 6: *Meloidogyne incognita* proteins for validation of putative effector.**

List of proteins in *M. incognita* proteome presenting at least one motif belonging to CLUMP5 (the most abundant in the proteome) and predicted to possess signal peptide and no transmembrane domain. Further details on the selection criteria can be found in the “Results & Discussion” section of the main text.

