## supplementary materials for "Identification and characterization of specific motifs in effector proteins of plant parasites using MOnSTER"

**­Supplementary Information**

**Configuration parameters for tools used in the study not in default modality.**

- cd-hit-2d with the following configuration: -s2 0 -c 0.90 -g1 -aL 0.30 -aS 0.30

s2, -c, -g parameter values are the default ones. -aL and -aS values are set so each sequence of a pair must cover at least 30 % of the other one.

- CD-HIT : s=0.8, c=0.99, g=1, aL=0.80, aS=0.80.
- STREME version 5.5.1, accessible at <https://meme-suite.org/meme/doc/download.html>.

Parameters used: **--minw 3 --maxw 5; --minw 3 --maxw 7**

- MERCI: <http://dtai-static.cs.kuleuven.be/ml/systems/MERCI/MERCI.zip>.

Parameter used: **-l 5 -fp 20; -l 7 -fp 20; -l 10 -fp 20**

**Supplementary figures**

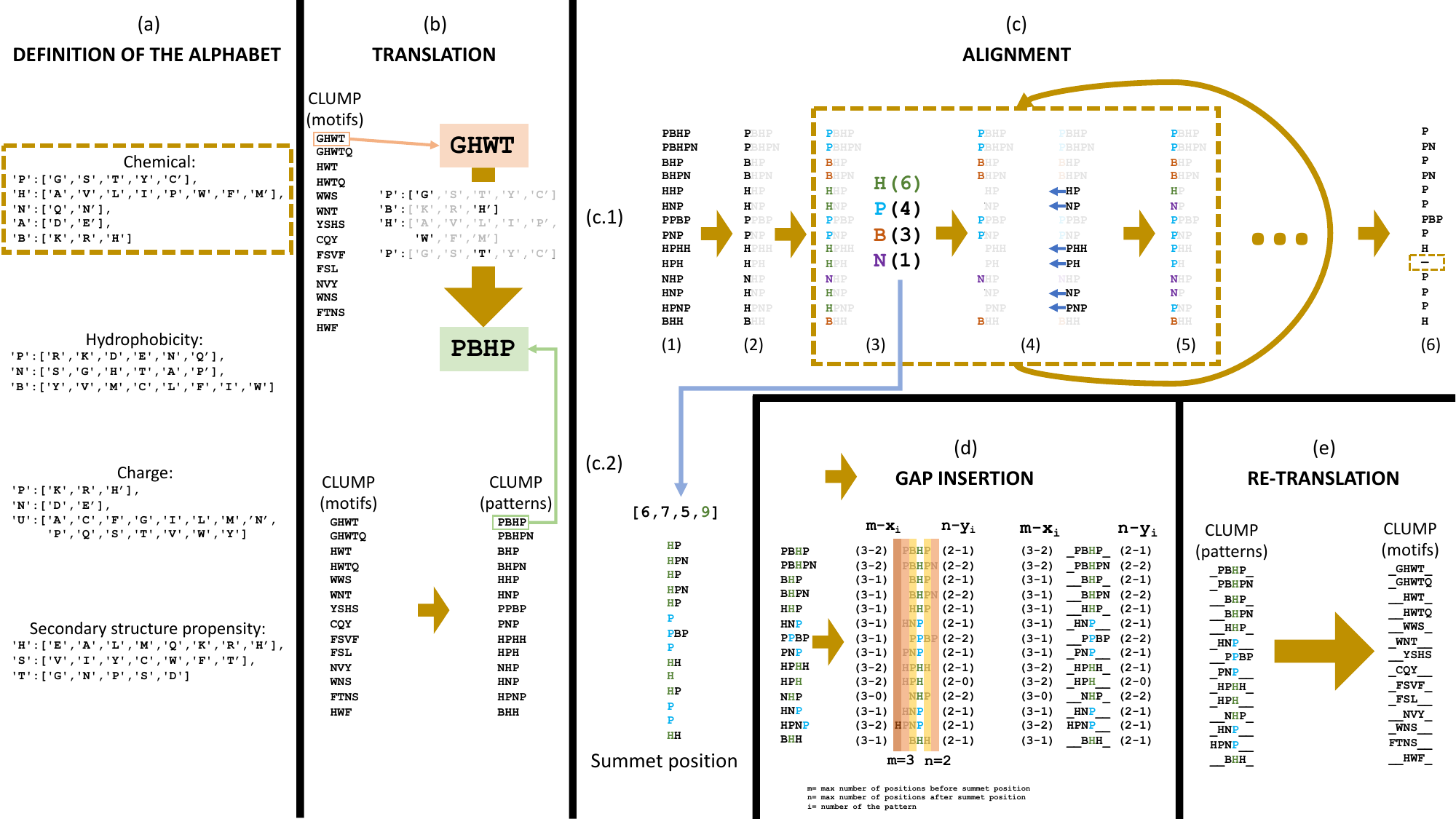

**Supplementary figure 1: PRO-MOCA algorithm.**

**(a) DEFINITION OF THE ALPHABET.** The user defines the alphabet to use to translate the protein sequences into patterns. Alphabet are constructed as pairs of keys, a letter defined by the user, and values, a list of amino acids to include in the group. Amino acids from the group will be replaced by the key in the sequence. PRO-MOCA already include the following alphabets. Chemical alphabet: ‘P’ = ’polar’, ‘H’ = hydrophobic, ‘N’= ‘neutral’, ‘A’ = ’acid’, ‘B’ = ‘basic’. Hydrophobicity alphabet: ‘P’ = ‘hydrophilic’, ’N’ = ‘neutral’, ‘B’ = ‘hydrophobic’. Charge alphabet: ‘P’ = ’positive’, ‘N’ = ’negative’, ’U’ = ‘neutral’. Secondary structure propensity alphabet: ‘H’ = ‘helix (α)’, ‘S’ = ‘sheet (β)’, ‘T’ = ‘turn’.

**(b) TRANSLATION.** Using the chosen alphabet (step (a)), the motifs in the CLUMP are translated from the protein alphabet (amino acids) into patterns of the given alphabet. For instance, using the ‘chemical’ alphabet, the ‘GHWT’ motif is translated into ‘PBHP’.

**(c) ALIGNMENT.** (c1) PRO-MOCA: (1) takes the list of patterns, (2) selects the first position of each pattern, (3) calculates how many times each letter is repeated at that position and finds the most repeated letter (PRO-MOCA stores this result at each iteration) (4) After that, patterns with the most repeated letter at the first position, lose that letter, and the rest of the pattern is shifted of -1 position. (5) The result is a list of patterns where part of the patterns has a new first position. From this point, the process (3)-(5) is repeated until (6) the result is a list where at least one pattern does not have any more letters. Until there is a pattern with no more letters (golden rectangle). (c2) Once the first part of the alignment is done and PRO-MOCA gets to (6), it finds the central position of the alignment. To do that, PRO-MOCA stored in (3) the number of repetitions of the most repeated letter. In (c2) PRO-MOCA calculates which is the highest value, and gets the corresponding list of patterns with those first positions.

(d) **GAP INSERTION**. The gap insertion is only allowed at the extremities of the patterns and allows to have an alignment where all the patterns have the same length. To do that, PRO-MOCA calculates, for each pattern, how many letters there are before (xi) and after (yi) the central position and calculates the maximum (m and n) of these values. After that, it subtracts m – xi and n – yi. The result of the subtraction is the number of gaps each pattern has before and after the central position.

**(e) RE-TRANSLATION.** In this step, PRO-MOCA re-translates the aligned patterns into aligned motifs

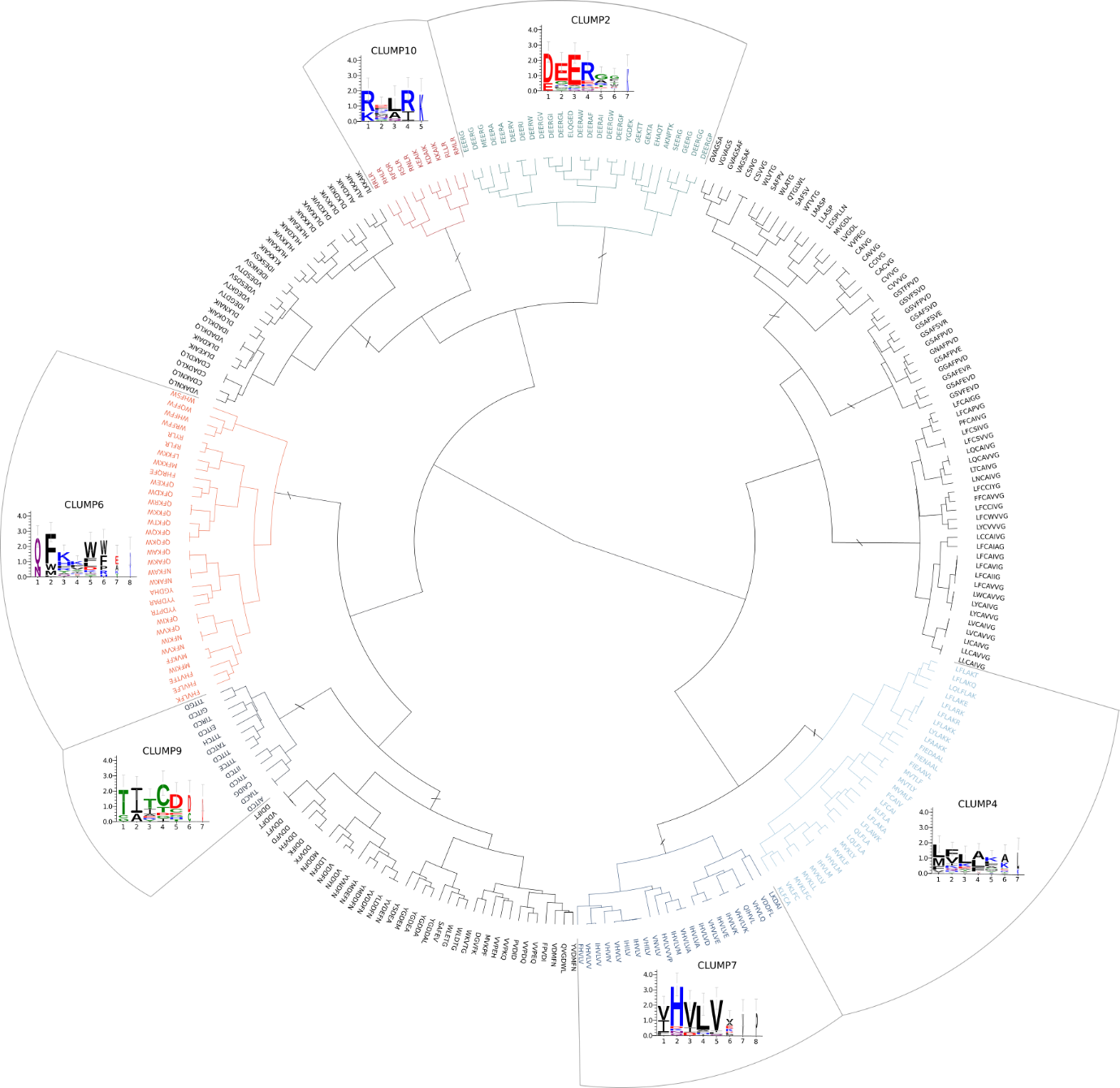

**Supplementary figure 2:** **Dendrogram of CLUMPs in Oomycetes.**

11 CLUMPs produced by MOnSTER (indicated with “/” sign). The coloured ones are those selected as best-scoring CLUMPs after MOnSTER-score calculation. Each best-scoring CLUMP is associated the motif logo; the alignment of the motif in each CLUMP is produced by PROMOCA and then WebLogo 3 is used to produce the image (the x-axis shows the AA position of the motif and the y-axis represents the log-transformed frequency of each AA in terms of bits of information).

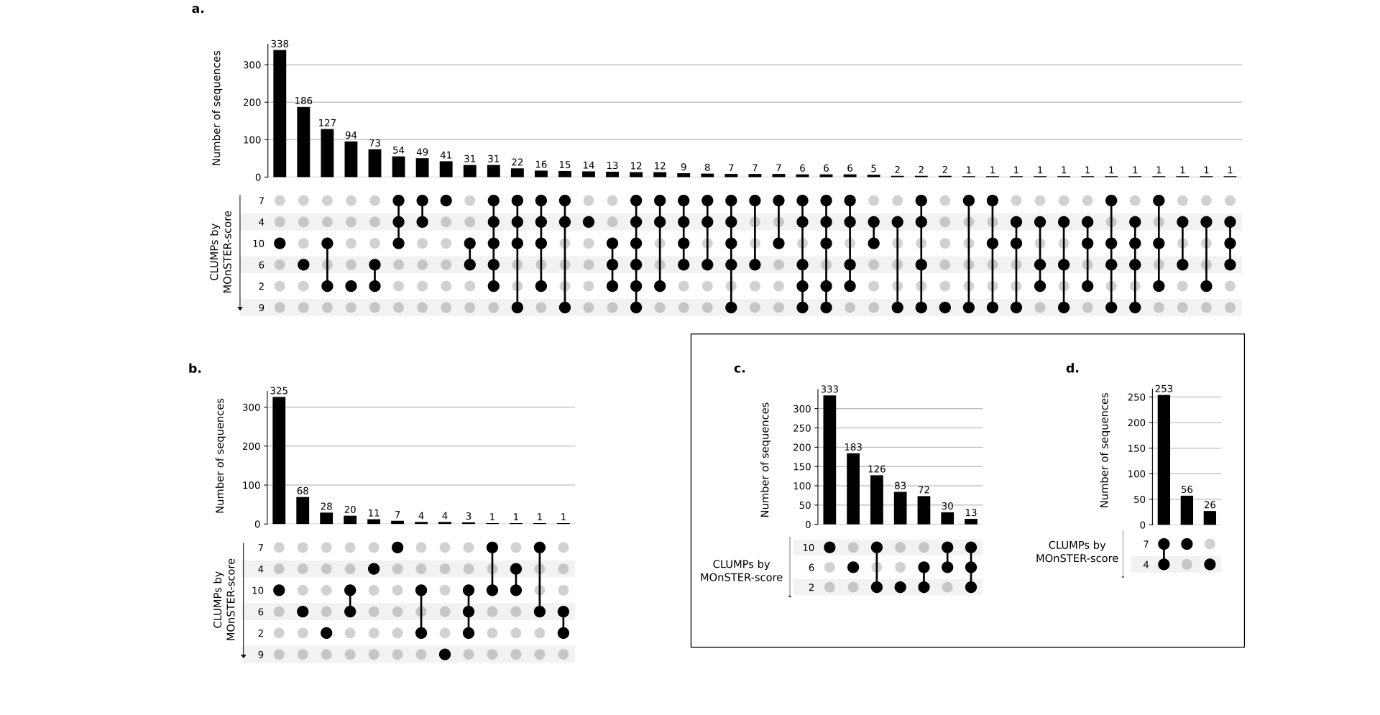

**Supplementary figure 3: Occurrences and co-occurrences of CLUMPs in positive and negative dataset – Oomycetes.**

Each upset plot represents the number of sequences in the respective dataset according to the presence of one or more CLUMPs ordered by MOnSTER-score. **(a)** positive dataset, **(b)** negative dataset, (c-d) specifically show the occurrence and co-occurrence of CLUMPs corresponding to RxLR and dEER (CLUMP10, 6, 2), and LxLFLAK-HVLVxxP (CLUMP7,4), in RxLR and Crinkler-effector sequences, respectively.

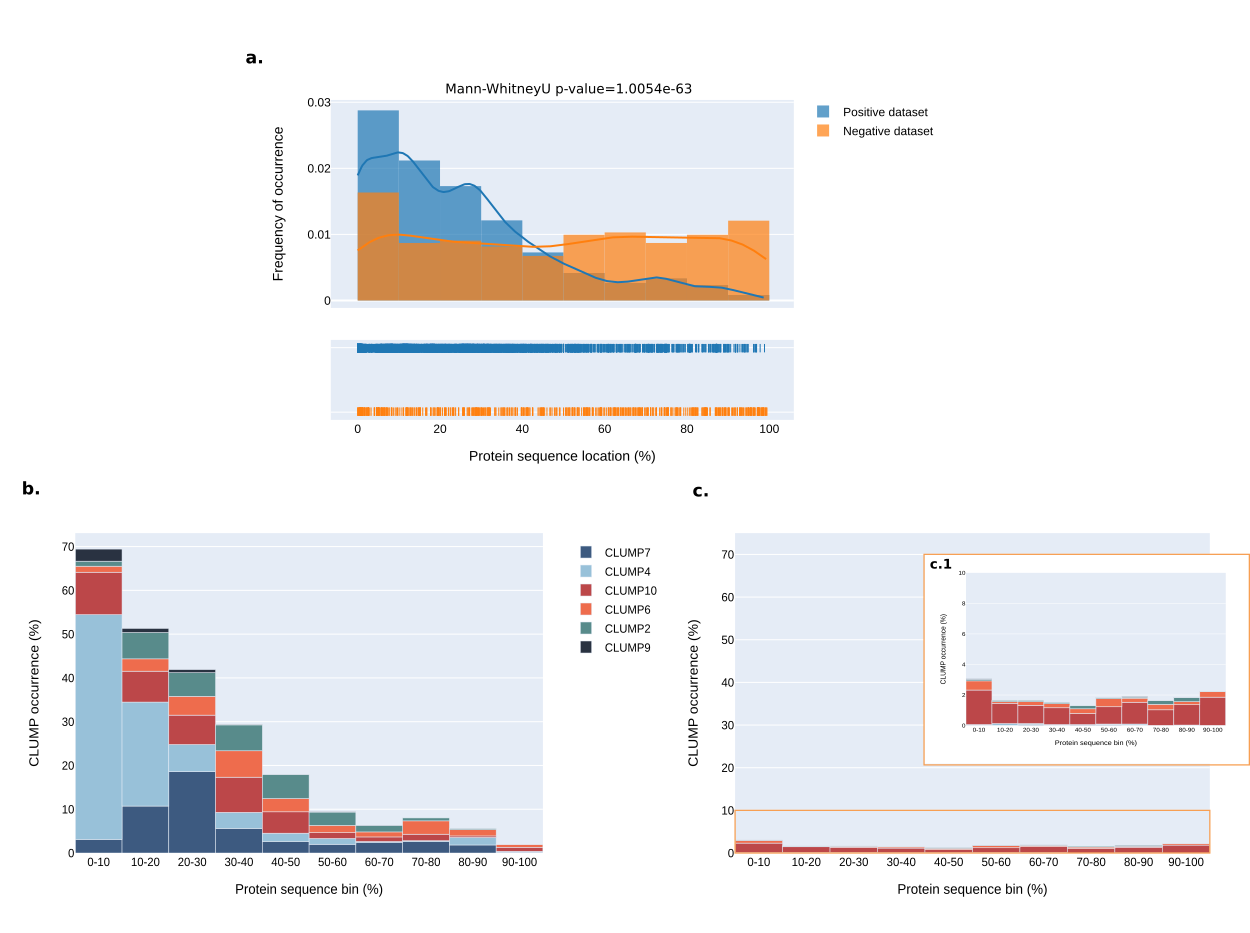

**Supplementary figure 4: Sequence position preference of motifs in CLUMPs – Oomycetes.**

**(a)** Shows the overall preference position of CLUMPs in the positive (blu trace) compared to the negative (orange trace) dataset. In the rug-plot each line corresponds to a CLUMP-motif position occurrence. **(b)** represents the same distribution in the positive dataset but divided by each CLUMP abundance in the sequence bin. **(c)** shows the same concept but in the negative dataset; **(c.1)** is a zoom-in of the original plot to better visualize the occurrences of CLUMPs. The distributions are all represented in bins of 10% each as absolute positions on the sequence.

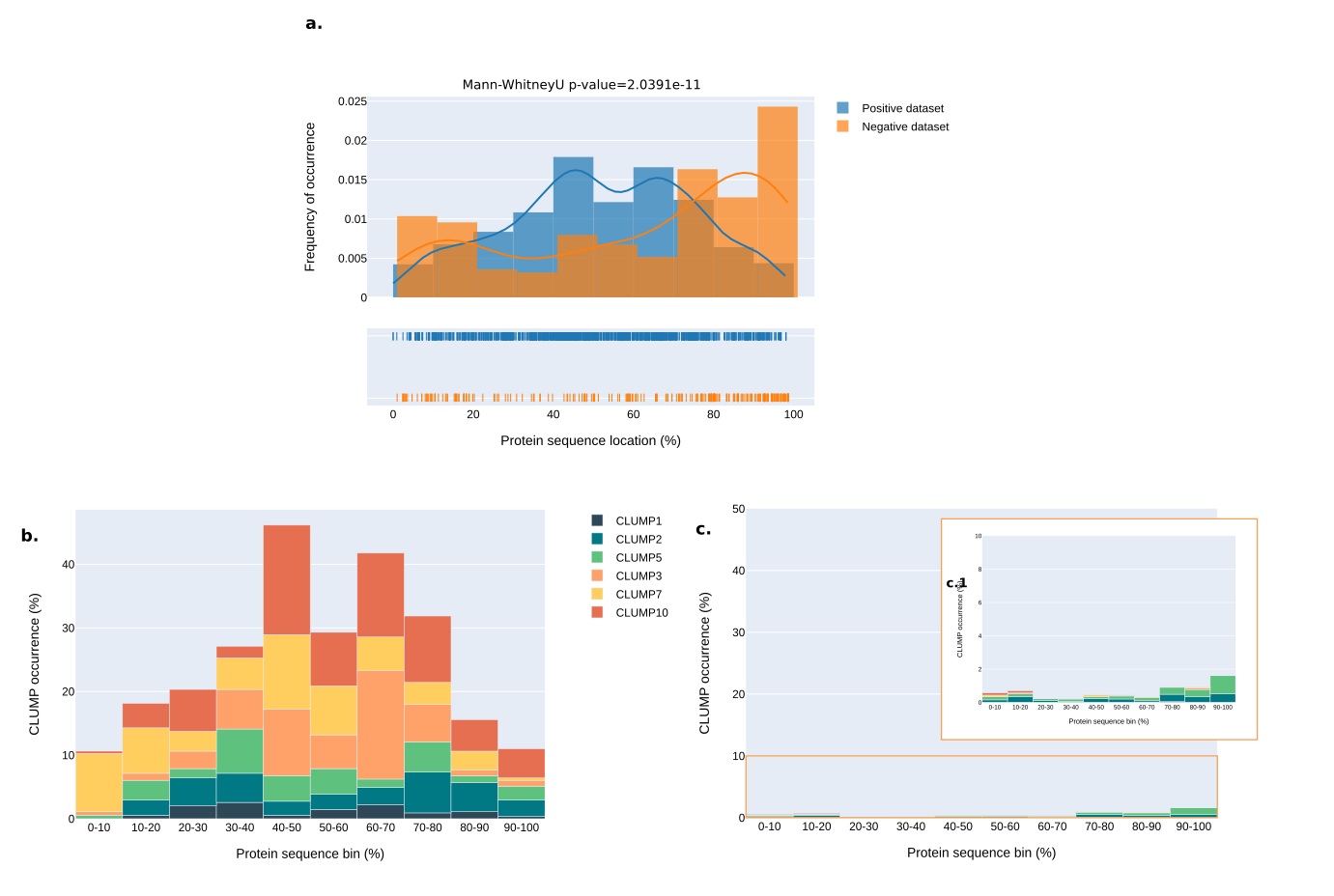

**Supplementary figure 5: Sequence position preference of motifs in CLUMPs – PPNs.**

**(a)** Shows the overall preference of CLUMPs in the positive (blu trace) compared to the negative (orange trace) dataset. In the rug-plot each line corresponds to a CLUMP-motif position occurrence. **(b)** represents the same distribution in the positive dataset but divided by each CLUMP abundance in the sequence bin. **(c)** shows the same concept but in the negative dataset; **(c.1)** is a zoom-in of the original plot to better visualize the occurrences of CLUMPs. The distributions are all represented in bins of 10% each as absolute positions on the sequence.

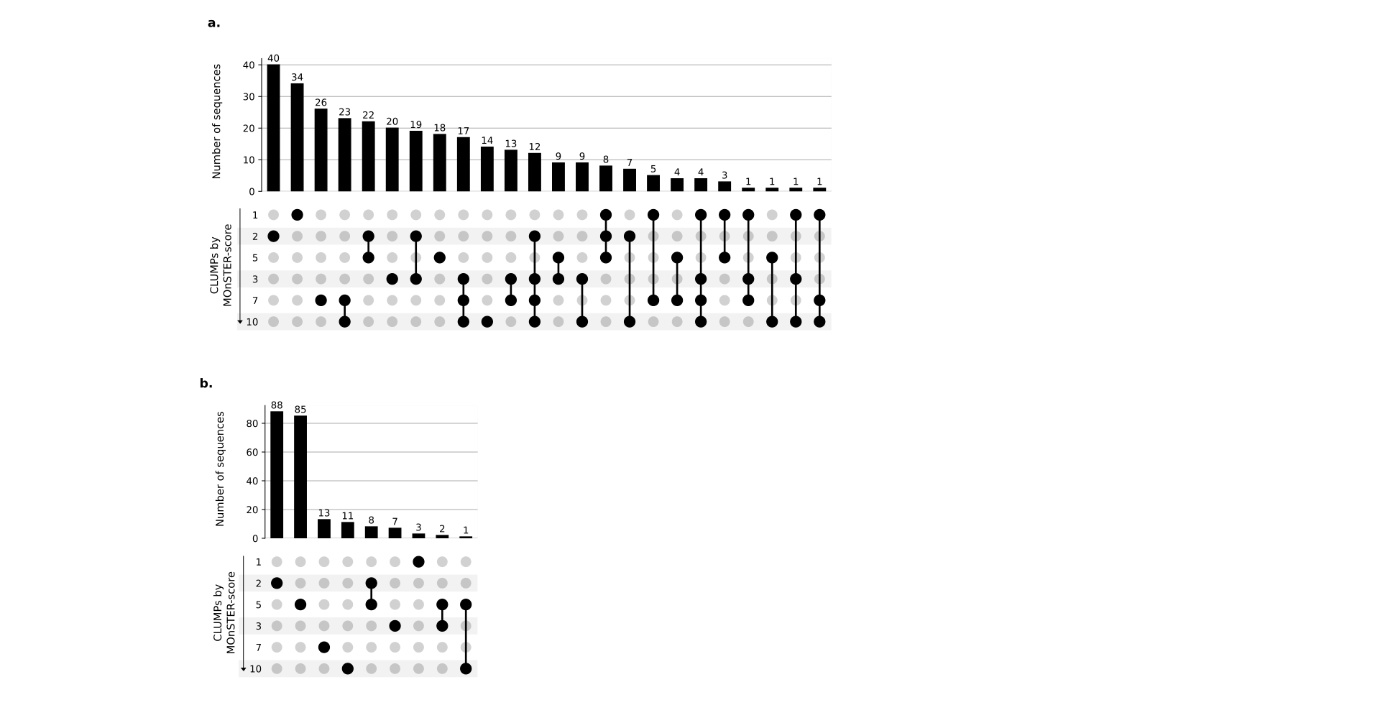

**Supplementary figure 6: Occurrence and co-occurrence of CLUMPs in PPNs.**

Representation of occurrence and co-occurrence of each best-scoring CLUMP in protein sequences, according to their MOnSTER-score. (a) shows the number of sequences and the occurrence of CLUMPs in the positive dataset, while (b) represents the same measures in the negative dataset.

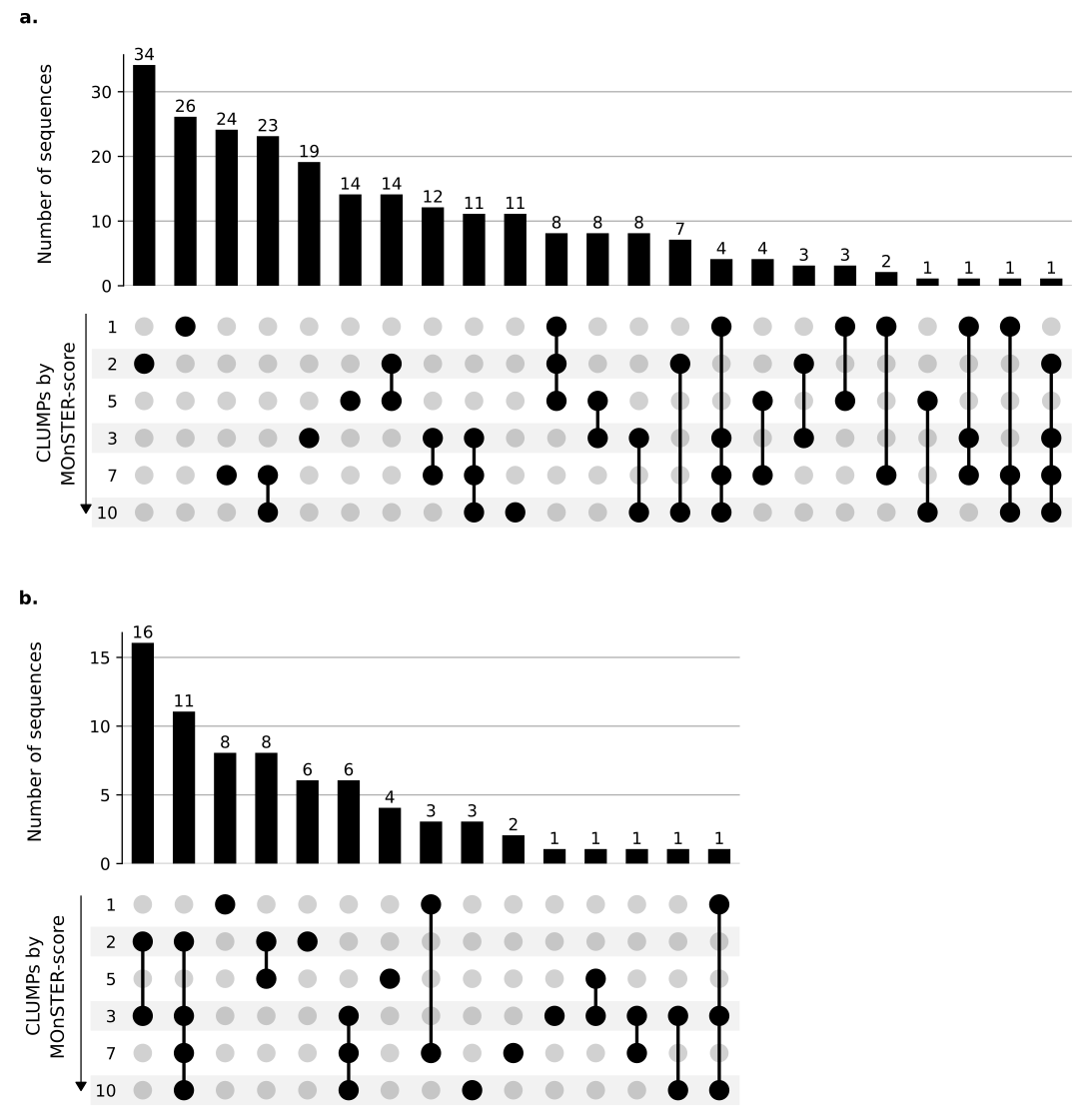

**Supplementary figure 7.** **Occurrence and co-occurrence of CLUMPs in proteins with or without the signal peptide.**

Representation of occurrence and co-occurrence of each best-scoring CLUMP in protein sequences, according to their MOnSTER-score. (a) shows the number of sequences and the occurrence of CLUMPs in the sequences bearing the signal peptide, while (b) represents the same measures in the sequences without the signal peptide.

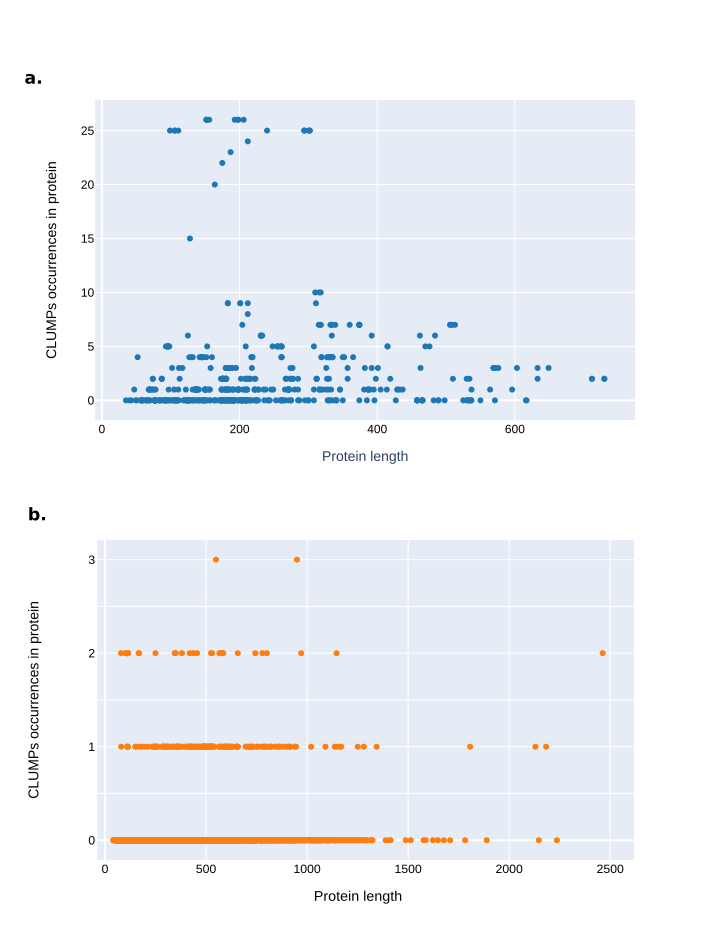

**Supplementary figure 8: Relation between CLUMPs abundance and protein sequence length – PPNs.**

**(a**) Shows a non-linear relationship between the occurrences of CLUMPs and the length of sequences in the positive dataset, while **(b)** represents the same but in the negative dataset.

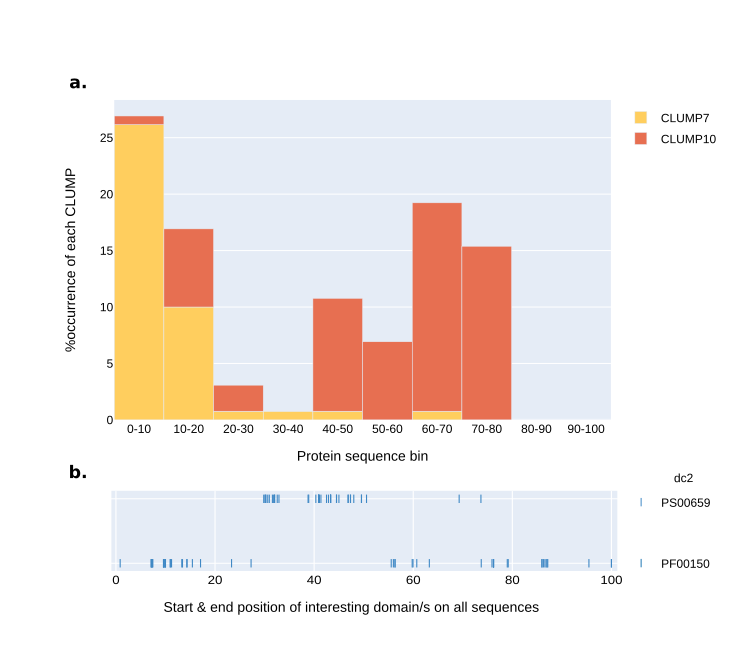

**Supplementary figure 9: Protein sequence position of motifs in CLUMP7 and 10 associated with glycosyl hydrolase family 5 domain (dc2).**

**(**a) CLUMP7 and CLUMP10 motifs-position and respective occurrence along selected protein sequences divided into bins representing 10% sequence portion. (**b**) rug-plot representing the start and end position of the domains (PS00659-ProSitePatterns and PF00150-Pfam) belonging to the class 2 of domains, namely glycosyl hydrolase family 5 domains.

**Supplementary table 1.1: Oomycetes’ positive dataset.**

Detailed description of the protein sequences in the oomycetes’ positive dataset including species, retrieving sources, and corresponding amino acid sequence.

**Supplementary table 1.2: Oomycetes’ negative dataset.**

Detailed description of the protein sequences in the oomycetes’ negative dataset including species, retrieving sources, and corresponding amino acid sequence.

**Supplementary table 2.1: PPNs’ positive dataset.**

Detailed description of the protein sequences in the PPNs’ positive dataset including species, retrieving sources, and corresponding amino acid sequence.

**Supplementary table 2.2: PPNs’ negative dataset.**

Detailed description of the protein sequences in the PPNs’ negative dataset including species, retrieving sources, and corresponding amino acid sequence.

**Supplementary table 3: Oomycetes’ MOnSTER-score.**

Summary table of MOnSTER-score for each CLUMP found in oomycetes dataset, ranked by higher score.

**Supplementary table 4: PPNs’ MOnSTER-score.**

Summary table of MOnSTER-score for each CLUMP found in PPNs dataset, ranked by higher score.

**Supplementary table 5: InterProScan domains and CLUMPs co-occurrences.**

Summary of co-occurrences of InterProScan domains in the six characteristic CLUMPs for PPN species.

**Supplementary table 6: *Meloidogyne incognita* proteins for validation of putative effector.**

List of proteins in *M. incognita* proteome presenting at least one motif belonging to CLUMP5 (the most abundant in the proteome) and predicted to possess signal peptide and no transmembrane domain.

Further details on the selection criteria can be found in the “Results & Discussion” section of the main text.
